# Genomic analysis reveals a functional role for myocardial trabeculae in adults

**DOI:** 10.1101/553651

**Authors:** Hannah V Meyer, Timothy JW Dawes, Marta Serrani, Wenjia Bai, Paweł Tokarczuk, Jiashen Cai, Antonio de Marvao, Daniel Rueckert, Paul M Matthews, Maria L Costantino, Ewan Birney, Stuart A Cook, Declan P O’Regan

## Abstract

Since being first described by Leonardo da Vinci in 1513 it has remained an enigma why the endocardial surfaces of the adult heart retain a complex network of muscular trabeculae – with their persistence thought to be a vestige of embryonic development. For causative physiological inference we harness population genomics, image-based intermediate phenotyping and *in silico* modelling to determine the effect of this complex cardiovascular trait on function. Using deep learning-based image analysis we identified genetic associations with trabecular complexity in 18,097 UK Biobank participants which were replicated in an independently measured cohort of 1,129 healthy adults. Genes in these associated regions are enriched for expression in the fetal heart or vasculature and implicate loci associated with haemodynamic phenotypes and developmental pathways. A causal relationship between increasing trabecular complexity and both ventricular performance and electrical activity are supported by complementary biomechanical simulations and Mendelian randomisation studies. These findings show that myocardial trabeculae are a previously-unrecognised determinant of cardiovascular physiology in adult humans.

## Main

The endocardial surfaces of the left and right ventricles are lined by a fenestrated network of muscular trabeculae which extend into the cavity. Their embryological development is driven by highly-conserved signalling pathways involving the endocardium-myocardium and extra-cellular matrix that regulate myocardial proliferation during cardiac morphogenesis.^1–6^ Cell lineage tracing suggests that trabeculae have a molecular and developmental identity which is distinct from the compact myocardium.^7^ The high surface area of trabeculae allows myocardial development to progress before the coronary circulation is established, and trabeculae are also vital to formation of the conduction system.^8, 9^ However, the role of trabeculae in the ventricular mechanics of healthy adults and why such a critical compartment of the cardiovascular system retains a complex inner surface is unclear. Theoretical analyses have shown that an absence of trabeculae requires greater strain to maintain cardiac output,^10^ and it has been proposed that their complex structure contributes to efficient intra-ventricular flow patterns.^11, 12^

Progress in understanding the functional role of organ components often relies on observing changes during physiological experiments or by identifying genes that selectively influence relevant morphology. Although capable of adaptation,^13–15^ myocardial trabeculation is an anatomical trait that does not have a clear stimulus-response relationship to investigate experimentally. Furthermore, complex pleiotropic effects of gene level lesions, such as knockouts, cause a complex array of abnormalities simultaneously affecting the trabeculae and other cardiovascular traits.^5^ Here we harness large scale population genomics, deep learning-driven image phenotyping and *in silico* modelling to determine causative effects of cardiovascular morphology on function. In a replicated genome-wide association study (GWAS) we identify loci associated with trabecular complexity and use Mendelian randomisation to provide a causal inference framework. We then use a finite-element model of the heart with physiologically-realistic loading conditions to determine the haemodynamic effect of varying trabecular complexity whilst keeping other parameters constant. This multi-disciplinary approach to physiological inference suggests a causal relationship between trabecular complexity and ventricular efficiency implicating loci associated with haemodynamic phenotypes and developmental pathways.

### Data overview

UK Biobank is a prospective cohort study collecting deep genetic and phenotypic data on approximately 500,000 individuals from across the United Kingdom, aged between 40 and 69 at recruitment, providing opportunities for the discovery of the genetic basis of complex traits.^16^ Of these, 100,000 participants are being recalled for enhanced phenotyping which includes cardiac magnetic resonance (CMR) imaging.^17^ Non-invasive data on a range of haemodynamic parameters are also collected at the time of imaging. Following automated image-quality control^18^ and exclusion of subjects with missing covariates, 18,097 unrelated participants that formed a well mixed population of European ethnicity (Figure 7 in the Supplement) were used for discovery (Table 1 and Figure 1D). An independent cohort comprising 1,129 healthy adults recruited in London, UK with both genotypes and CMR imaging was used for validation.^19^

**Table 1.** Participant characteristics. Measurements depicted in mean ± standard deviation. End-systolic volume, end-diastolic volume and mass are indexed to body surface area. BP, Blood pressure; FD, fractal dimension.

| Characteristics | UK Biobank | UK Digital Heart Study |
| --- | --- | --- |
| Age (years) | 55 ± 7 | 41 ± 13 |
| Body surface area (m<sup>2</sup>) | 1.90 ± 0.21 | 1.85 ± 0.20 |
| Sex | 9402/8694 (m/f) | 621/508 (m/f) |
| Haemodynamics |  |  |
| Systolic BP (mmHg) | 137 ± 18 | 120 ± 14 |
| Diastolic BP (mmHg) | 81 ± 10 | 79 ± 9 |
| Heart rate (beats/minute) | 62 ± 10 | 62 ± 21 |
| Cardiac output (l/min) | 5.5 ± 1.3 | 6.0 ± 2.0 |
| Left ventricular volumetry |  |  |
| Ejection fraction (%) | 59.5 ± 6.0 | 65.3 ± 6.9 |
| End-systolic volume (ml/m<sup>2</sup>) | 31.9 ± 8.6 | 28.2 ± 7.9 |
| End-diastolic volume (ml/m<sup>2</sup>) | 78.3 ± 13.9 | 81.2 ± 14.4 |
| Mean global FD | 1.169 ± 0.028 | 1.215 ± 0.038 |

**Figure 1.**
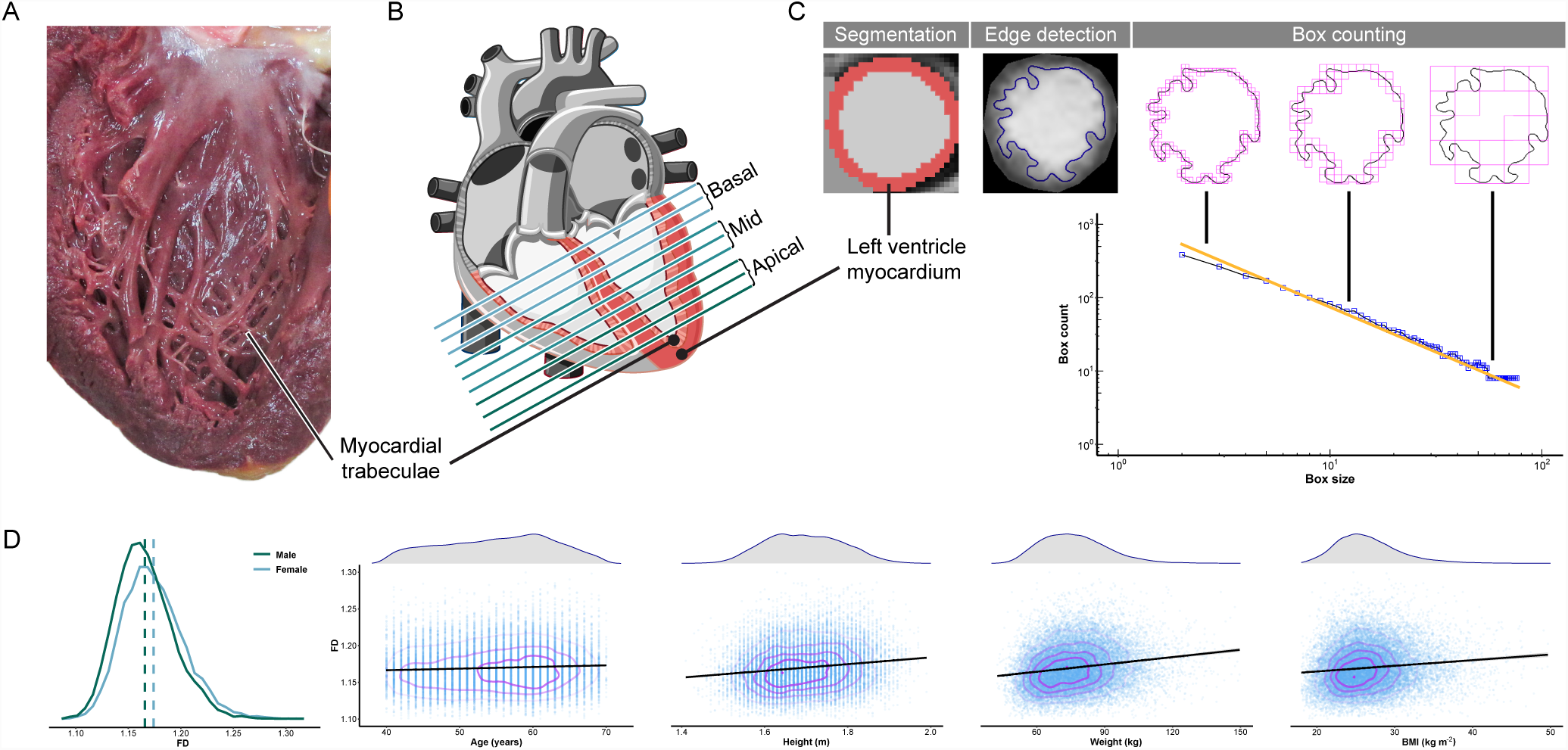
Trabeculation phenotypes and co-variates. A) Macroscopic cut pathological section of the left ventricle demonstrating the fenestrated network of muscular trabeculae lining the endocardial surface. B) Diagram of the heart illustrating the positioning of sections acquired during cardiac magnetic resonance (CMR) imaging for the assessment of trabecular complexity. C) Deep learning image segmentation was used for anatomical annotation of each pixel in the CMR dataset and to define an outer region of interest for subsequent fractal analysis. A binary mask was taken of the image followed by edge detection of the trabeculae. Box-counting across a range of sizes generated a log-log plot from which the gradient of a least-squares linear regression defined the fractal dimension. D) Distribution of fractal dimension (FD) and its relation to co-variates used in the association study.

### Fractal dimension analysis of left ventricular trabeculation

We used a fully convolutional network for automated left ventricular segmentation and volumetry.^20^ The complexity of myocardial trabeculae (Figure 1A) was quantified by the scale-invariant ratio of fractal dimension (FD, Figure 1C).^21^ To account for variations in cardiac size and for consistent anatomical comparisons within and between populations we interpolated the data to 9 slices (see Figure 6 in the Supplement) which were equally divided into basal, mid-ventricular and apical thirds (Figure 1B). An identical analytic pipeline was performed in the validation cohort.

### Genome-wide association analysis of fractal dimension

We performed a linear model for genetic association of 14,180,594 genetic variants on each of the 9 interpolated slice FD measures of 18,097 individuals using anthropometric variables as co-variates (Figure 1D). These genome-wide association studies showed low inflation and many individual loci passing the commonly used genome-wide association threshold of 5 × 10^*-*8^ after adjustment for multiple testing by the effective number of tests (*T*_*e f f*_ = 6.6; see Figure 9 and Figure 10 in the Supplement). We performed a meta-analysis over all 9 slices to both gain power and have a single discovery process. Figure 2B shows the resulting 16 independent loci from this meta-analysis and the individual slice which the loci are associated with. Four loci were only discovered using this joint meta-analysis approach (Figure 2B, orange circles); the remaining 12 loci have distinct patterns of association across the different slices but show a biologically-plausible variation of effect size from base to apex (Figure 11 in the Supplement). Representative myocardial borders associated with a locus on chromosome 8 (see section below) are depicted in Figure 2D. An analogous association study including end-diastolic volume as a co-variate is shown in Figure 12 in the Supplement. Both studies lead to the discovery of the same loci, indicating that FD associations are independent of ventricular size. To replicate our findings we analysed the independent cohort of 1,129 people, applied the same image analysis pipeline and ran an equivalent genetic association study (10,673,171 genetic variants). As expected given the lower sample size, few of the associations passed a genome-wide threshold, however nearly all the estimates of effect direction were concordant (91% of comparisons concordant) between the two studies (correlation of beta estimates: *r*^2^ = 0.516, Figure 13 in the Supplement). Permutation tests generating empirical concordance distributions show that the observed concordance is extremely unlikely to be observed by chance (*p*_empirical_ < 10^*-*5^).

**Figure 2.**
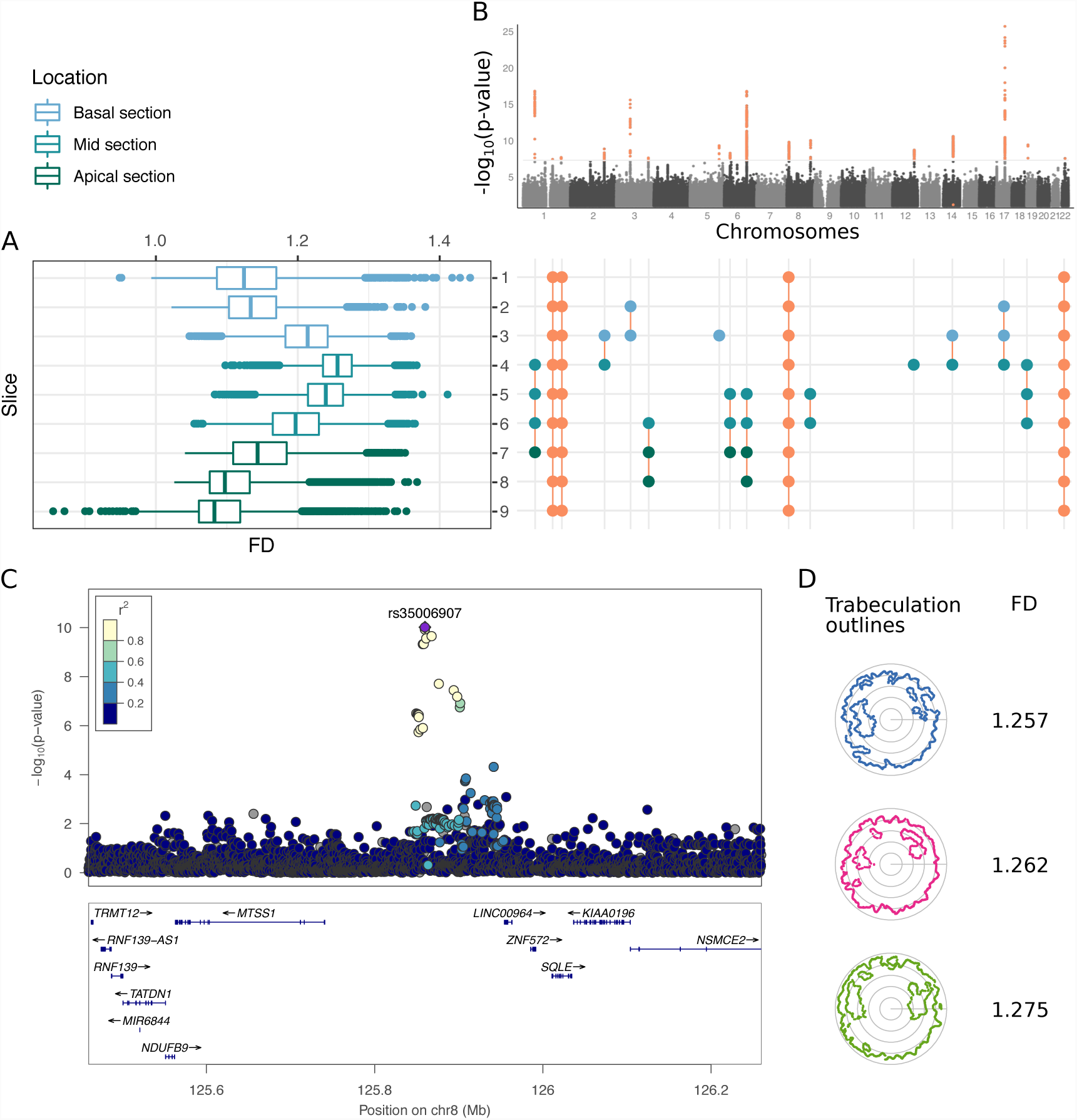
Genetic associations of left ventricular trabeculation. A. Box-plots of FD measurements per slice, colour-coded by cardiac region. B. Manhattan plot of meta-analysis p-values, with loci passing the genome-wide significance threshold 5 × 10^*-*8^ highlighted in orange (top). Diagram showing the slices driving the genetic association signal (compare Figure 9 in the Supplement): circles indicate a locus being associated with respective slice and region. Loci marked in orange circles have no individual association 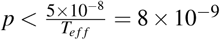 (where *T*_*e f f*_ = 6.6 is the effect number of independent phenotypic tests) and were only discovered in the meta-analysis. C) Locus zoom of the locus on chromosome 8, associated with slices 5 and 6. D) Registered, trabecular outlines at slice 5 representing the median FD for individuals with homozygous major (blue), heterozygous (pink) and homozygous minor (green) genotype for rs35006907.

### Functional and molecular associations of the discovered loci

We systematically analysed the 16 discovered loci with the rich genetic resources of other studies, drawing from both the extensive GWAS Catalog,^22^ and more recent phenome-wide associations (PheWAS) from UK Biobank.^23^ Table 2 summarises our findings (for details on loci see Table 4 in the Supplement).

**Table 2.**
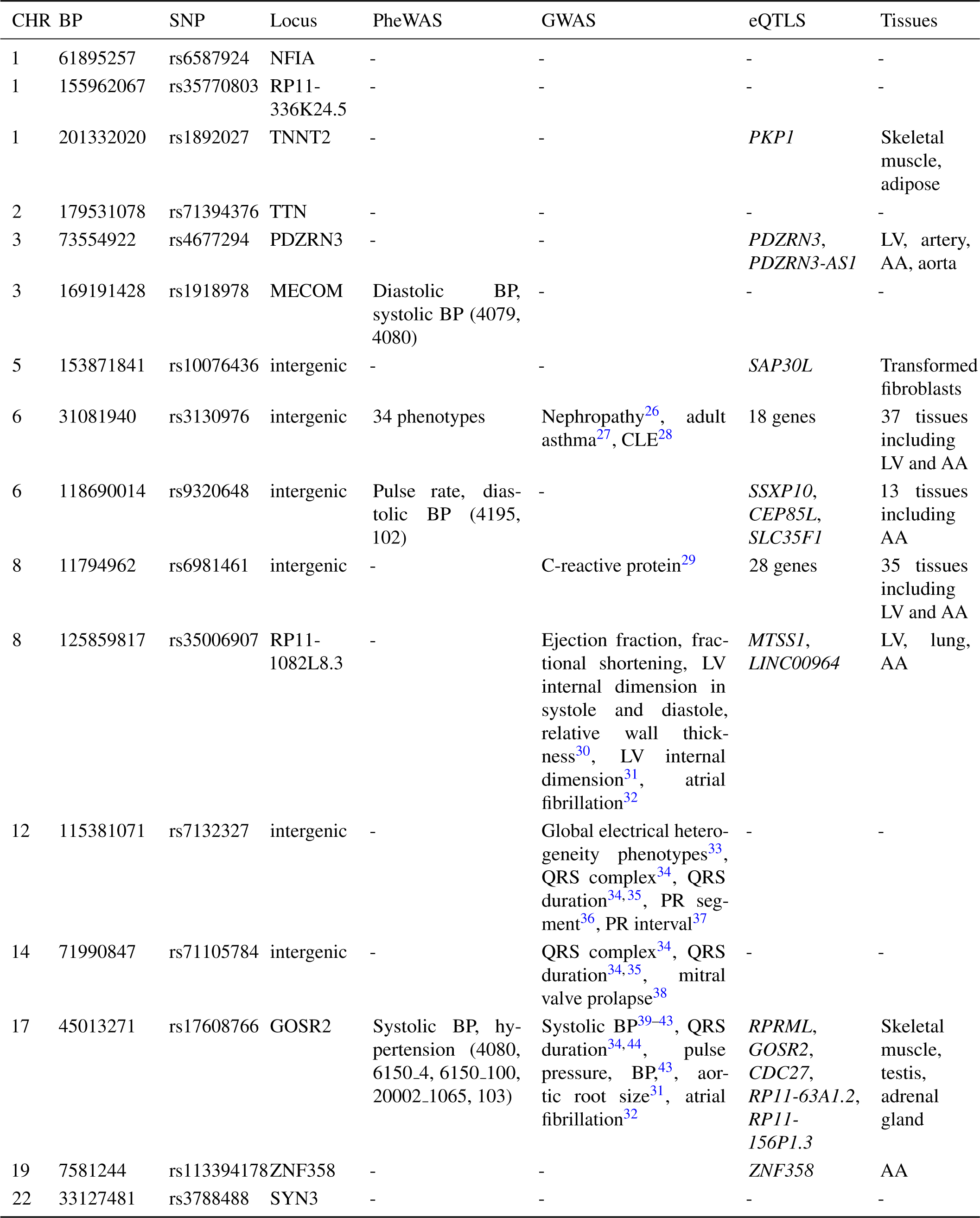
Annotations of trabeculation-associated loci. Overview of the 16 independent loci discovered in the trabeculation GWAS. The genetic variant with the lowest p-value per locus is shown. Annotations: PheWAS: phase 2 PheWAS described by^24^ (phenotype reference ID in parentheses), GWAS: GWAS catalogue^22^, eQTLs and tissues: GTEx catalogue v7^25^. Chromosomes (CHR), base pair positions (BP) and Locus: GRCh37 (Ensembl GRCh37 Release 95). No entry indicated by -. BP, Blood pressure; LV, left ventricle; AA, atrial appendage.

**Table 3.** Finite element modelling. Steady-state measurements of the left ventricle, selectively varying only trabecular complexity, at constant total myocardial mass with identical boundary conditions.

| Haemodynamics | Low FD | High FD |
| --- | --- | --- |
| Peak systolic pressure (mmHg) | 93 | 123.6 |
| End-systolic pressure (mmHg) | 88 | 119.8 |
| Diastolic pressure (mmHg) | 50 | 65.9 |
| Left ventricular volumes and function |  |  |
| End-diastolic volume (ml) | 114.8 | 160.4 |
| End-systolic volume (ml) | 54.8 | 80.1 |
| Stroke volume (ml) | 60 | 80.3 |
| Stroke work (J) | 0.62 | 1.09 |
| Cardiac output (l/min) | 3.6 | 4.8 |
| Contractility (mmHg/ml) | 1.61 | 1.50 |
| Mass (g) | 97.7 | 97.7 |
| Fractal dimension | 1.11 | 1.8 |

**Table 4.** Characteristics of trabeculation-associated loci. Overview of the 16 independent loci discovered in the trabeculation GWAS. The genetic variant with the lowest p-value per locus is shown. Chromosomes (CHR), base pair positions (BP), ID, Locus and Type based on GRCh37 (Ensembl GRCh37 Release 95). Allele frequency (AF), frequency describing allele (A 0) and alternative allele (A 1) based on discovery cohort data. INFO denotes the imputation quality score of impute2^73^. No entry indicated by -.

| CHR | SNP | BP | A_0 | A_1 | AF | INFO | P | Ensemble ID | Locus | Type |
| --- | --- | --- | --- | --- | --- | --- | --- | --- | --- | --- |
| 1 | rs6587924 | 61895257 | C | A | 0.51 | 1.00 | 1.57E-17 | ENSG00000162599 | NFIA | intron |
| 1 | rs35770803 | 155962067 | G | GT | 0.63 | 0.96 | 3.71E-08 | ENSG00000224276 | RP11-336K24.5 | intron |
| 1 | rs1892027 | 201332020 | T | C | 0.71 | 1.00 | 2.01E-08 | ENSG00000118194 | TNNT2 | intron |
| 2 | rs71394376 | 179531078 | A | AATGT | 0.21 | 1.00 | 1.40E-09 | ENSG00000155657 | TTN | intron |
| 3 | rs4677294 | 73554922 | T | A | 0.36 | 0.99 | 2.61E-16 | ENSG00000121440 | PDZRN3 | intron |
| 3 | rs1918978 | 169191428 | A | G | 0.58 | 0.98 | 2.40E-08 | ENSG00000085276 | MECOM | intron |
| 5 | rs10076436 | 153871841 | C | G | 0.36 | 1.00 | 4.90E-10 | - | intergenic | - |
| 6 | rs3130976 | 31081940 | T | C | 0.29 | 1.00 | 5.29E-09 | - | intergenic | - |
| 6 | rs9320648 | 118690014 | A | C | 0.58 | 1.00 | 1.67E-17 | - | intergenic | - |
| 8 | rs6981461 | 11794962 | T | C | 0.44 | 0.98 | 1.64E-10 | - | intergenic | - |
| 8 | rs35006907 | 125859817 | C | A | 0.31 | 0.99 | 9.55E-11 | ENSG00000255080 | RP11-1082L8.3 | intron |
| 12 | rs7132327 | 115381071 | T | C | 0.27 | 1.00 | 1.97E-09 | - | intergenic | - |
| 14 | rs71105784 | 71990847 | C | CCTGT | 0.25 | 1.00 | 2.63E-11 | - | intergenic | - |
| 17 | rs17608766 | 45013271 | T | C | 0.15 | 1.00 | 1.98E-26 | ENSG00000108433 | GOSR2 | 3' UTR<br>intron |
| 19 | rs113394178 | 7581244 | C | A | 0.61 | 0.96 | 3.81E-10 | ENSG00000198816 | ZNF358 | intron |
| 22 | rs3788488 | 33127481 | T | C | 0.27 | 0.99 | 2.81E-08 | ENSG00000185666 | SYN3 | intron |

Ten of the 16 loci are also associated with at least one component of heart function, such as blood pressure, pulse rate, QRS duration, left ventricular structure and function (for details see Table 5 and Table 6 in the Supplement). Broadly the more apical slices (6, 7, 8) have an overlap with blood pressure phenotypes, whereas the basal to mid-ventricular slices (3, 4) have QRS complex associated phenotypes.

**Table 5.**
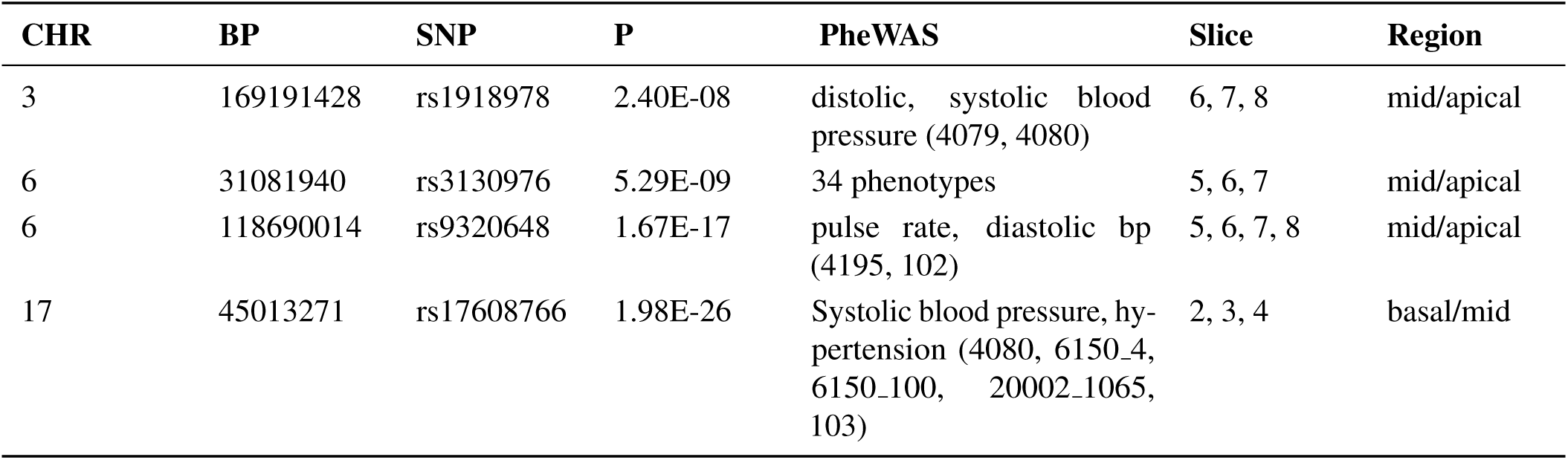
Loci annotation in UK Biobank PheWAS. Based on the phase 2 PheWAS described by^24^. The phenotype reference ID are given in parentheses). No entry indicated by -.

| CHR | BP | SNP | P | PheWAS | Slice | Region |
| --- | --- | --- | --- | --- | --- | --- |
| 3 | 169191428 | rs1918978 | 2.40E-08 | diastolic, systolic blood pressure (4079, 4080) | 6, 7, 8 | mid/apical |
| 6 | 31081940 | rs3130976 | 5.29E-09 | 34 phenotypes | 5, 6, 7 | mid/apical |
| 6 | 118690014 | rs9320648 | 1.67E-17 | pulse rate, diastolic bp (4195, 102) | 5, 6, 7, 8 | mid/apical |
| 17 | 45013271 | rs17608766 | 1.98E-26 | Systolic blood pressure, hypertension (4080, 6150_4, 6150_100, 20002_1065, 103) | 2, 3, 4 | basal/mid |

**Table 6.** Loci annotation in published GWAS. Based on entries in the GWAS catalogue^22^. No entry indicated by -. BP, Blood pressure; LV, left ventricle/left ventricular; AA, atrial appendage; CLE, Cutaneous lupus erythematosus.

| CHR | BP | SNP | P | GWAS | Slice | Region |
| --- | --- | --- | --- | --- | --- | --- |
| 6 | 31081940 | rs3130976 | 5.29E-09 | Nephropathy<sup>26</sup>, adult asthma<sup>27</sup>, CLE<sup>28</sup> | 5, 6, 7 | mid/apical |
| 8 | 11794962 | rs6981461 | 1.64E-10 | C-reactive protein<sup>29</sup> | multi-trait | - |
| 8 | 125859817 | rs35006907 | 9.55E-11 | Ejection fraction, fractional shortening, LV internal dimension in systole and diastole, relative wall thickness<sup>30</sup>, LV internal dimension<sup>31</sup>, atrial fibrillation<sup>32</sup> | 5, 6 | mid |
| 12 | 115381071 | rs7132327 | 1.97E-09 | Global electrical heterogeneity phenotypes<sup>33</sup>, QRS complex<sup>34</sup>, QRS duration<sup>34,35</sup>, PR segment<sup>36</sup>, PR interval<sup>37</sup> | 4 | mid |
| 14 | 71990847 | rs71105784 | 2.63E-11 | QRS complex<sup>34</sup>, QRS duration<sup>34,35</sup>, mitral valve prolapse<sup>38</sup> | 3, 4 | basal/mid |
| 17 | 45013271 | rs17608766 | 1.98E-26 | Systolic blood pressure<sup>39-43</sup>, QRS duration<sup>34,44</sup>, pulse and blood pressure<sup>43</sup>, aortic root size<sup>31</sup>, atrial fibrillation<sup>32</sup> | 2, 3, 4 | basal/mid |

We compared our loci to the extensive GTEx catalog^25^ of gene expression quantitative trait loci (eQTL; Table 2 and Table 7 in the Supplement). Nine of the 16 loci showed an overlap with a GTEx locus; in eight cases at least one of the eQTL tissue was either cardiac tissue or skeletal muscle; in one case the only significant tissue was transformed fibroblasts. A particularly strongly annotated association is on chromosome 8, in a region of open chromatin that is an expression QTL for the *MTSS1* gene (Figure 2C). This locus is also associated to a variety of cardiac structure and function phenotypes (Table 2, rs35006907), and the lead genetic variant located is in a region of open chromatin in heart tissues (ENSR00000868700, ENSEMBL regulatory build^45^).

**Table 7.** Loci annotation in GTEx. Based on the GTEx catalogue v7^25^. No entry indicated by -. BP, Blood pressure; LV, left ventricle; AA, atrial appendage.

| CHR | BP | SNP | P | gtex - eQTLs | gtex - tissues | Slice | Region |
| --- | --- | --- | --- | --- | --- | --- | --- |
| 1 | 201332020 | rs1892027 | 2.01E-08 | *PKP1* | Skeletal muscle, adipose | multi-trait | - |
| 3 | 73554922 | rs4677294 | 2.61E-16 | *PDZRN3*,<br>*PDZRN3-AS1* | LV, artery, AA, aorta | 2, 3 | basal |
| 5 | 153871841 | rs10076436 | 4.90E-10 | *SAP30L* | Transformed fibroblasts | 3 | basal |
| 6 | 31081940 | rs3130976 | 5.29E-09 | 18 genes | 37 tissues including LV and AA | 5, 6, 7 | mid/apical |
| 6 | 118690014 | rs9320648 | 1.67E-17 | *SSXP10*, *CEP85L*,<br>*SLC35F1* | 13 tissues including AA | 5, 6, 7, 8 | mid/apical |
| 8 | 11794962 | rs6981461 | 1.64E-10 | 28 genes | 35 tissues including LV and AA | multi-trait | - |
| 8 | 125859817 | rs35006907 | 9.55E-11 | *MTSS1*,<br>*LINC00964* | LV, lung, AA | 5, 6 | mid |
| 17 | 45013271 | rs17608766 | 1.98E-26 | *RPRML*, *GOSR2*,<br>*CDC27*, *RP11-*<br>*63A1.2*, *RP11-*<br>*156P1.3* | Skeletal muscle, testis, adrenal gland | 2, 3, 4 | basal/mid |
| 19 | 7581244 | rs113394178 | 3.81E-10 | *ZNF358* | AA | 4, 5, 6 | mid |

In addition to previously reported associations, we were interested in the functional annotations of our GWAS results. As the per-slice GWAS (Figure 2B) suggested regionally-driven signals, we conducted genome-wide associations of basal (slices 1-3), mid-ventricular (slices 4-6) and apical (slices 7-9) FD. We analysed all genetic variants of these association results for enrichment in regulatory and functional annotations, using GARFIELD.^46^ GARFIELD accounts for LD structure and local gene density and derives the statistical significance of the functional enrichment (odds ratios) by fitting a logistic regression model. The strongest associations of the genetic loci were to open-chromatin regions in fetal heart tissue, particularly in the mid and apical regions (Figure 3, Figure 14 in the Supplement).

**Figure 3.**
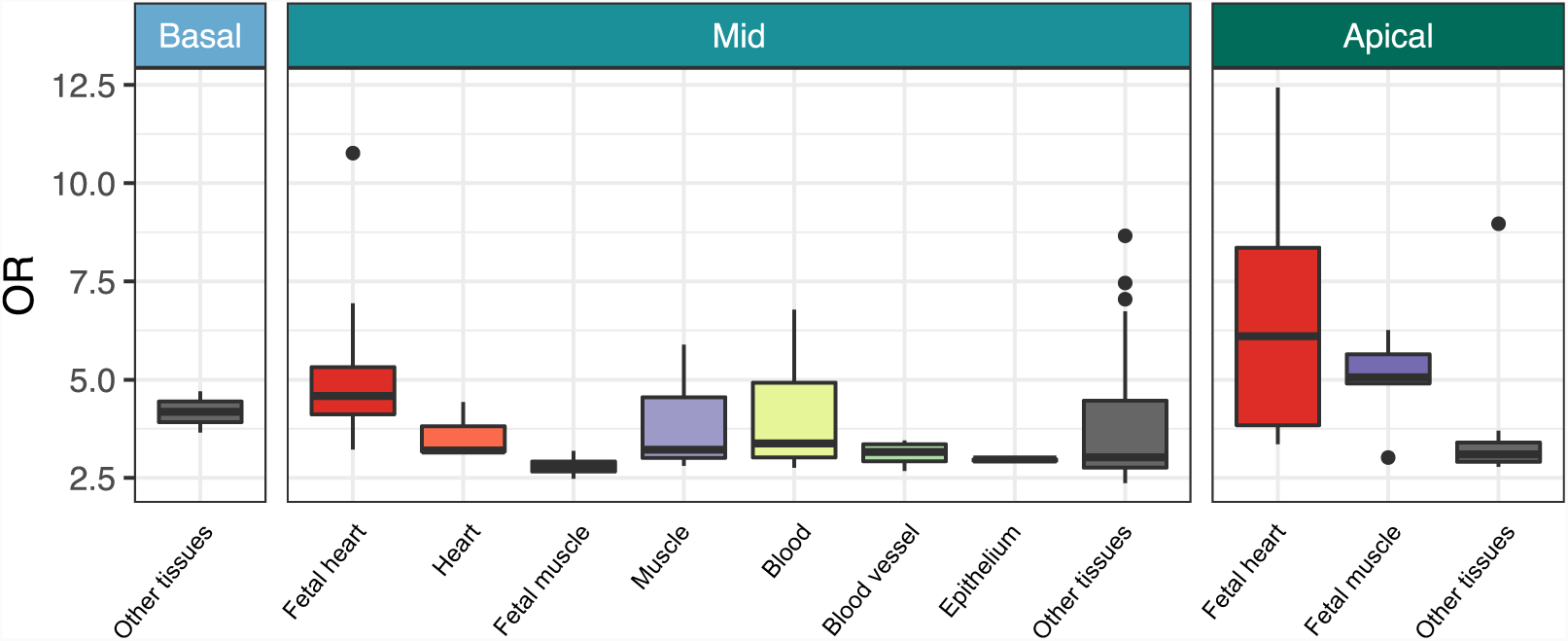
Enrichment of trabeculation associated variants in DNaseI Hypersensitive sites. GARFIELD was used to compute the functional enrichment (odds ratio, OR) of genetic variants associated with the trabeculation phenotypes ((*p* < 10^*-*6^) for open chromatin regions. Enrichment analyses were done for each cardiac region independently and results across all available studies per tissue are depicted in box-plots. ‘Other tissues’ comprises the collection of all available DNaseI Hypersensitive datasets within the GARFIELD datasets.

Overall the discovered loci are mainly linked with either molecular or physiological cardiac phenotypes, but with a variety of specific associations. Some loci are likely developmental, such as the locus on chromosome 8 associated with *MTSS1*, affecting many aspects of cardiac function whereas other loci have more specific associations. Amongst the well-annontated loci electrical physiology is a common theme, for example rs17608766 which is also associated with QRS duration, blood pressure, cardiac anatomy and expression QTLs to three genes in skeletal muscle.

### Causal effects of trabeculae on function

A visualisation of cardiac mechanics during systole and diastole is provided by plotting a closed loop describing the relationship between left ventricular pressure and left ventricular volume at multiple time points during a complete cardiac cycle (Figure 4A).^47^ This also enables quantification of stroke work and contractility which are aspects of ventricular function inaccessible by other methods.^48^ To understand how trabeculae influence cardiac function we therefore assessed the relationship between FD and pressure-volume parameters of the left ventricle both in human populations and *in silico* models. We performed this in UK Biobank participants by analysing non-invasive estimates of central pressures combined with volumetric CMR data. In parallel, we developed a cardiovascular simulation, using finite element analysis of the left ventricle in a haemodynamic circuit, to determine the effect of selectively varying trabecular complexity on cardiac performance. In UK Biobank participants, increasing FD was associated with higher stroke volume (*β* = 0.52), stroke work (*β* = 0.67) and cardiac index (*β* = 0.12), findings which were replicated in the biomechanical simulation and suggest a causal relationship between trabecular complexity and these haemodynamic responses (Figure 4 and Table 3).

**Figure 4.**
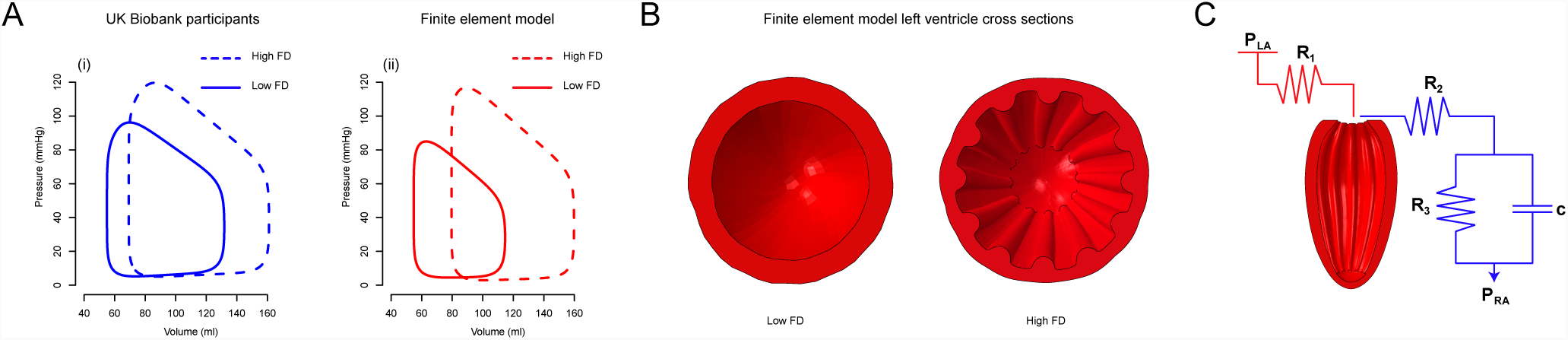
Relationship between trabecular complexity and cardiac function. A) The relationship between left ventricular volume and pressure during a cardiac cycle is represented by a closed loop where its width represents stroke volume, and its enclosed area is stroke work, ie the work done by the ventricle to eject a volume of blood. The variation in pressure-volume relationship with respect to trabecular fractal dimension in i) UK Biobank participants and ii) *in silico* biomechanical modelling. B) Mid short axis cross sections of the finite element model of the left ventricle, looking towards the apex, at different trabecular complexities. C) The ventricular model was in series with pre-load (red) and after-load circuits (blue) defining left atrial pressure (P_*LA*_), right atrial pressure (P_*RA*_), inflow resistance (R_1_), aortic resistance (R_2_), and both peripheral resistance (R_3_) and vascular capacitance (c).

To test the hypothesis that trabecular complexity is causally-related to increased cardiac performance in humans we used a two-sample Mendelian randomisation framework^49^ with the discovered independent loci as instrumental variables which are not subject to the inevitable confounding present in observational studies.^50^ We used FD as our exposure variable and stroke volume as the outcome, where the outcome associations were derived from an independent, UK Biobank discovery GWAS.^49^ A variety of Mendelian randomisation techniques exist, differing in their assumptions about pleiotropy of the instrumental variables, measurement uncertainty of exposure and outcome variables and cohort independence (for details refer to Mendelian randomisation in the Supplement). We tested a number of different techniques, each addressing different assumptions and found parameter estimates that robustly support a causal relationship between increasing trabecular complexity and stroke volume in the mid to apical regions of the left ventricle (Figure 5A; Figure 16 and Tables 8 to 10 in the Supplement). The directionality of the relationship was confirmed by assessing the difference in the genetic correlations of exposure and outcome (MR Steiger^51^, Table 12 in the Supplement). Estimates of the F-statistic indicate no weak instrument bias despite potential sample overlap between these UK Biobank studies (Table 13 in the Supplement). We did not observe large pleiotropic effects for the genetics variants associated with trabeculation and stroke volume (Table 11 in the Supplement).

**Table 8.** Mendelian randomisation results for FD associations in basal region. MR results are shown for outcome variables and respective IDs (outcome) derived from MR base^49^. nsnp specifies the number of snps present in both the exposure (FD associations in the basal region) and outcome study. b and se are the causal effect size estimate and standard error of the study and MR method, pval the p-pvalue of the MR analysis. For MR Egger, the magnitude of dilution bias is *I*^2^ = 0.987639, i.e. an approximately 1.3% underestimation of effect size is expected. F statistics and their lower bound can be found in Table 13. For details on methods and statistics refer to Mendelian randomisation.

| outcome | method | nsnp | b | se | pval |
| --- | --- | --- | --- | --- | --- |
| QRS duration - id:UKB-b:2240 | MR Egger | 4 | 1.715434244 | 0.94 | 0.209293571 |
| QRS duration - id:UKB-b:2240 | Inverse variance weighted | 4 | 0.083285136 | 0.33 | 0.799811846 |
| QRS duration - id:UKB-b:2240 | Weighted median | 4 | 0.328557247 | 0.1 | 0.001265225 |
| QRS duration - id:UKB-b:2240 | Weighted mode | 4 | 0.332062678 | 0.1 | 0.048254772 |
| LV stroke volume - id:UKB-b:6025 | MR Egger | 4 | 0.731133019 | 0.4 | 0.205440047 |
| LV stroke volume - id:UKB-b:6025 | Inverse variance weighted | 4 | 0.166408988 | 0.1 | 0.111546601 |
| LV stroke volume - id:UKB-b:6025 | Weighted median | 4 | 0.215216828 | 0.12 | 0.079654046 |
| LV stroke volume - id:UKB-b:6025 | Weighted mode | 4 | 0.251075291 | 0.16 | 0.210188394 |

**Table 9.** Mendelian randomisation results for FD associations in mid-ventricular region. MR results are shown for outcome variables and respective IDs (outcome) derived from MR base^49^. nsnp specifies the number of snps present in both the exposure (FD associations in the mid-ventricular region) and outcome study. b and se are the causal effect size estimate and standard error of the study and MR method, pval the p-pvalue of the MR analysis. For MR Egger, the magnitude of dilution bias is *I*^2^ = 0.9793806, i.e. an approximately 2.1% underestimation of effect size is expected. F statistics and their lower bound can be found in Table 13. For details on methods and statistics refer to Mendelian randomisation.

| outcome | method | nsnp | b | se | pval |
| --- | --- | --- | --- | --- | --- |
| QRS duration - id:UKB-b:2240 | MR Egger | 10 | 2.553323002 | 0.81 | 0.013572352 |
| QRS duration - id:UKB-b:2240 | Inverse variance weighted | 10 | 0.126581324 | 0.18 | 0.480315964 |
| QRS duration - id:UKB-b:2240 | Weighted median | 10 | 0.306828562 | 0.11 | 0.004693126 |
| QRS duration - id:UKB-b:2240 | Weighted mode | 10 | 0.368601288 | 0.14 | 0.024087935 |
| LV stroke volume - id:UKB-b:6025 | MR Egger | 10 | 1.516849049 | 0.54 | 0.022444816 |
| LV stroke volume - id:UKB-b:6025 | Inverse variance weighted | 10 | 0.372633253 | 0.1 | 0.000269965 |
| LV stroke volume - id:UKB-b:6025 | Weighted median | 10 | 0.380108679 | 0.12 | 0.001709506 |
| LV stroke volume - id:UKB-b:6025 | Weighted mode | 10 | 0.403724703 | 0.17 | 0.042085665 |

**Table 10.**
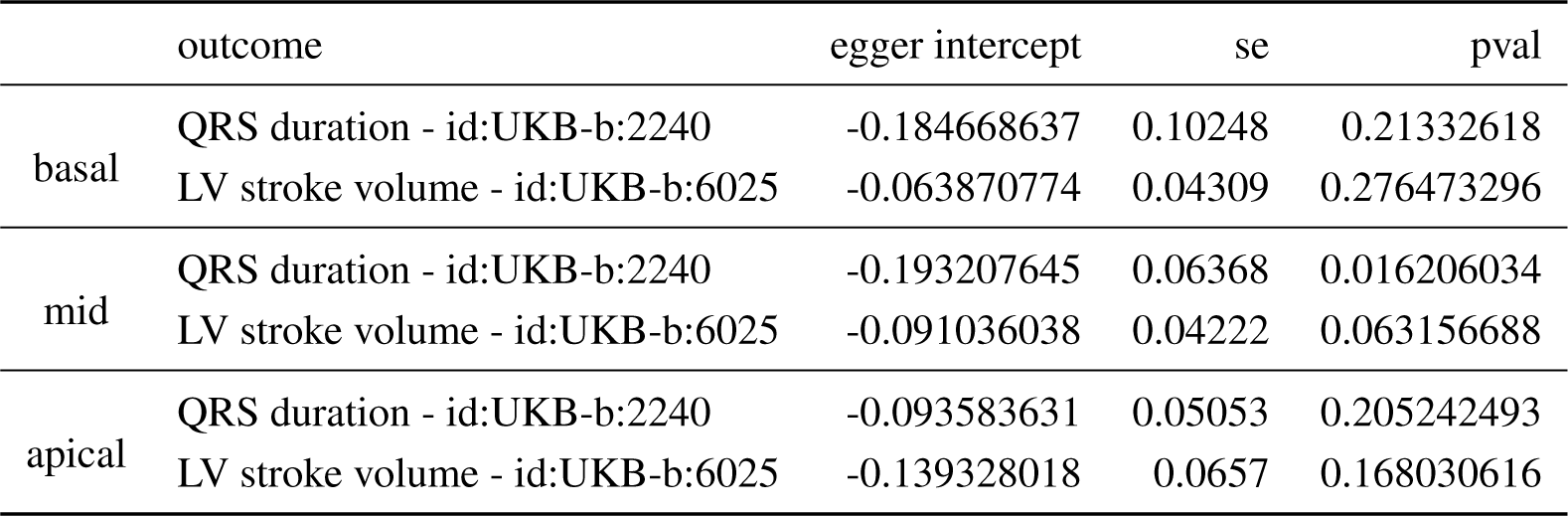
Mendelian randomisation results for FD associations in apical region. MR results are shown for outcome variables and respective IDs (outcome) derived from MR base^49^. nsnp specifies the number of snps present in both the exposure (FD associations in the apical region) and outcome study. b and se are the causal effect size estimate and standard error of the study and MR method, pval the p-pvalue of the MR analysis. For MR Egger, the magnitude of dilution bias is *I*^2^ = 0.9804846, i.e. an approximately 2% underestimation of effect size is expected. F statistics and their lower bound can be found in Table 13. For details on methods and statistics refer to Mendelian randomisation.

**Table 11.**
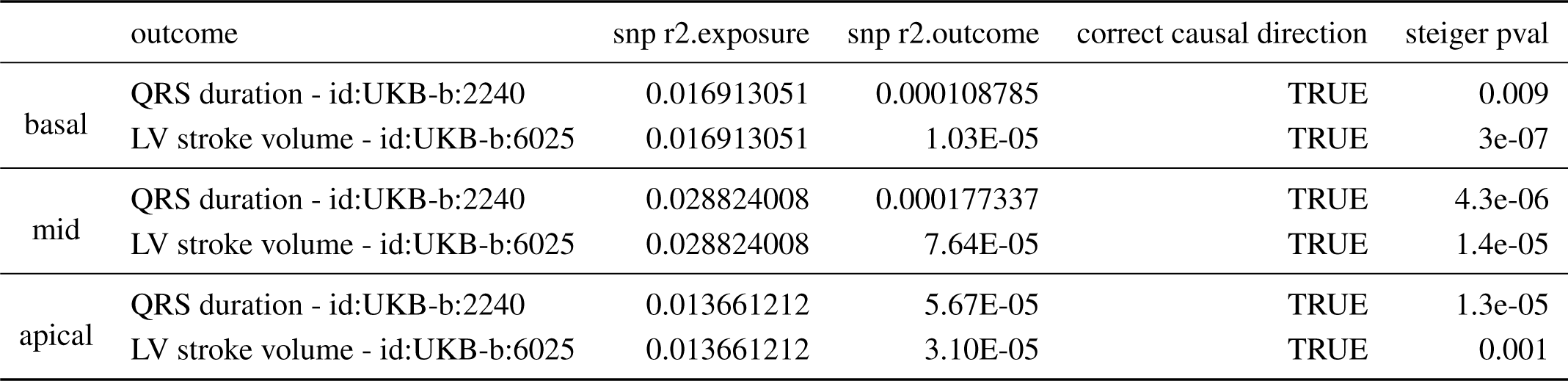
Pleiotropy assessment with MR-Egger for FD associations. MR Egger results are shown for outcome variables and respective IDs (outcome) derived from MR base^49^. egger intercept and se are the pleiotropy estimate and its standard error, pval the p-pvalue of the MR egger intercept. For details on MR Egger refer to Mendelian randomisation.

**Table 12.**
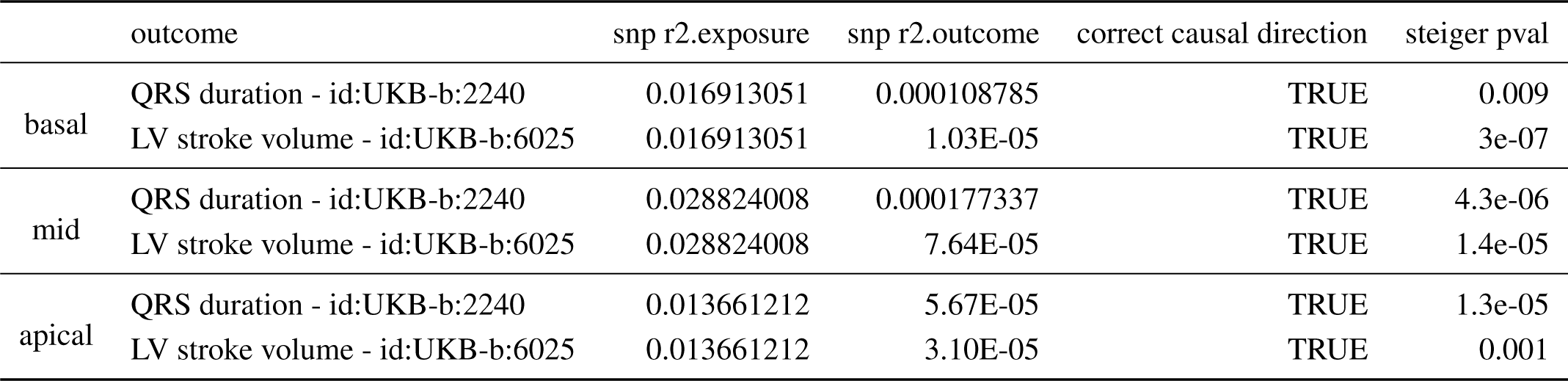
Steiger directionality analysis for causality on MR analyses. MR Steiger directionality results are shown for outcome variables and respective IDs (outcome) derived from MR base^49^. snp r2.exposure and snp r2.outcome are the *r*^2^ estimates of the correlation of all instrumental variables with the exposure and outcome traits, respectively. snp r2.exposure differ dependent on how many genetic variants overlapped between the FD associations and the outcome study - correlation only estimated for variants present in outcome and exposure. correct causal direction specifies if the hypothesised direction of FD upstream of the outcome trait is true. steiger pval specified the p-value of the MR Steiger test. For details on MR Steiger refer to Mendelian randomisation.

**Table 13.** F statistic of MR studies. The F statistic depends on the sample size in the exposure study (N exposure), number of IVs (IV), and the proportion of variance in the risk factor explained by the IVs (r2). It is computed by 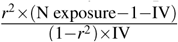. As the F statistic is a cohort estimate of the unknown population F parameter a lower bound of the F parameter was estimated according to [95, Appendix A3]. All lower bounds are *≥* 10, so despite possible sample overlap between the the UKB studies (id:UKB-b:2240, id:UKB-b:6025) and the FD GWAS, no strong weak instrument bias would be expected Mendelian randomisation.

|  | outcome | IV | N exposure | N outcome | r2 exposure | F statistic | Lower bound F |
| --- | --- | --- | --- | --- | --- | --- | --- |
| basal | QRS duration - id:UKB-b:2240 | 4 | 18097 | 463013 | 0.0169 | 77.81 | 63.74 |
|  | LV stroke volume - id:UKB-b:6025 | 4 | 18097 | 463013 | 0.0169 | 77.81 | 63.74 |
| mid | QRS duration - id:UKB-b:2240 | 10 | 18097 | 463013 | 0.0288 | 53.68 | 46.22 |
|  | LV stroke volume - id:UKB-b:6025 | 10 | 18097 | 463013 | 0.0288 | 53.68 | 46.22 |
| apical | QRS duration - id:UKB-b:2240 | 4 | 18097 | 463013 | 0.0137 | 62.65 | 50.08 |
|  | LV stroke volume - id:UKB-b:6025 | 4 | 18097 | 463013 | 0.0137 | 62.65 | 50.08 |

**Figure 5.**
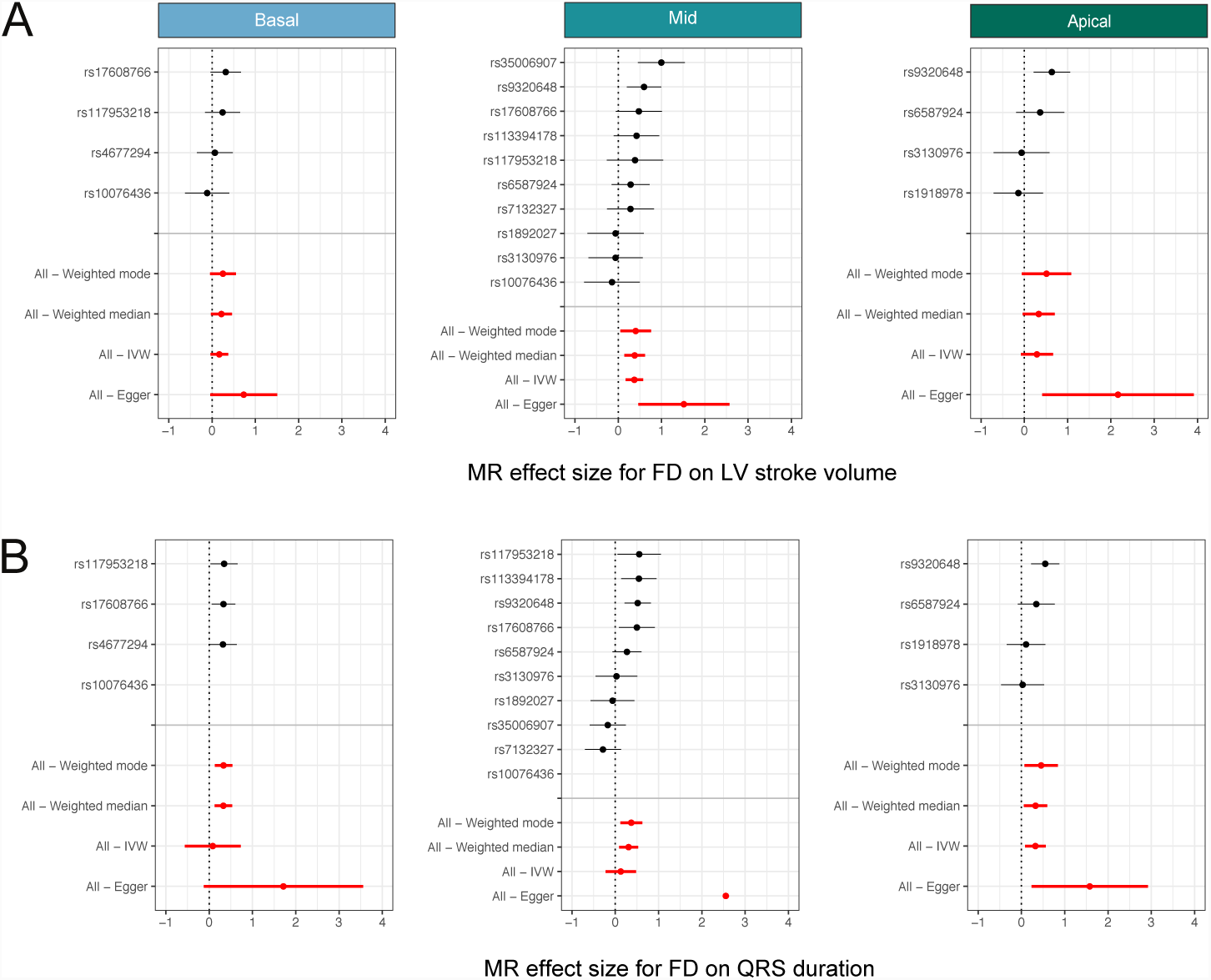
Mendelian randomisation analysis. Forest plots depicting the contribution of each genetic variant to the overall estimate (black) and combined as a single genetic instrument (red) for the four tested MR methods. A) Effect of trabeculation on stroke volume. B) Effect of trabeculation on QRS duration.

Trabeculae give rise to the ventricular conduction system during embryonic heart development,^9^ and we found a positive correlation in UK Biobank of an electrical phenotype (QRS duration) with FD (Figure 15 in the Supplement). We tested this hypothesis using the Mendelian randomisation approach described above and see a consistent positive effect of FD on QRS duration (Figure 5B; Figure 17 and Tables 8 to 10 and 12 in the Supplement) - confirming a mechanistic link between complex structural morphology and conduction characteristics in the left ventricle.^52^

## Discussion

Here we use a multi-disciplinary approach for physiological inference to discover causal relationships between trabecular complexity and both ventricular efficiency and conduction traits - implicating 16 loci associated with cardiovascular phenotypes and developmental pathways.

Trabeculae were first described by anatomists in the 16th Century,^53^ but beyond a role in maintenance of blood flow during early stages of cardiac morphogenesis (before expansion of the compact zone) their significance in adults has been unclear.^54^ Using deep learning for large scale precision cardiac phenotyping enabled us to perform the first reported GWAS of trabecular morphology. We found associations with trabecular complexity in loci related to cardiac function phenotypes, gene expression variation in cardiac tissues and cardiac development chromatin annotation that were independent of biophysical variables and ventricular volume. For example, the locus associated with the *MTSS1* gene supports a role for cytoskeletal signaling in trabecular morphology extending previously-reported associations with myocardial geometry.^31^ The bundle branches and peripheral ventricular conduction system develop from sub-endocardial myocytes of trabeculae,^9^ and we discovered three loci (rs7132327, rs71105784, rs17608766) previously associated with electrical phenotypes indicating the relevance of trabecular characteristics to conduction traits in adults. In addition, a number of loci overlap with well-established cardiac genes (*TNT, TNNT2*), linked to sarcomeric function and cardiac morphogeneis, that are related to a spectrum of hyper-trabeculation phenotypes.^55–57^ As well as multiple loci associated with cardiovascular traits, this study has also robustly implicated a number of loci which have yet to be associated with known phenotypes - whether physiological or disease-related - pointing to previously unexplored pathways related to trabecular formation.

For causal inference on the role of trabeculae on myocardial function we used complementary finite-element modelling of the left ventricle and Mendelian randomisation using trabeculation-associated loci. Our *in silico* simulations indicated a significant effect of trabecular complexity on ventricular performance, allowing for a higher end-diastolic volume and consequently a higher stroke volume to be achieved at the same atrial pressure. A concordant relationship between trabecular complexity and ventricular performance was observed using non-invasive clinical observations indicating that this association also holds for human populations. This hypothesis was supported by a variety of Mendelian randomisation techniques, addressing different model assumptions, taking the discovered genetic variants as a collection of instrumental variables that perturb trabeculation independently of environmental confounders. These inferential analyses show strong support for causal relationships between trabecular complexity and both functional and conduction phenotypes. The triangulation of theoretical models, observational data and genomics^58^ is persuasive evidence that trabeculae are not simply vestigial features of development but are powerful determinants of cardiac performance in adults. Taken with strong enrichment in cardiovascular tissues, these findings provide a foundation for exploring new structural and biological mechanisms that sustain heart function.

## Methods

All analyses in this study can be found here: https://github.com/ImperialCollegeLondon/fractalgenetics/.

### Phenotyping

#### Participants

For UK Biobank, approximately 500,000 community-dwelling participants aged 40–69 years were recruited across the United Kingdom between 2006 and 2010.^23^ Baseline summary characteristics of the cohort can be viewed on the UK Biobank data showcase (http://www.ukbiobank.ac.uk). Since 2014, a subset of participants are being recalled for CMR who live in proximity to the facility in Stockport, England, UK and 19,701 consecutive participants were included in our analysis. All subjects provided written informed consent for participation in the study, which was also approved by the National Research Ethics Service (11/NW/0382). Our study was conducted under terms of access approval number 18545. The validation cohort was drawn from the UK Digital Heart Project - a single-centre prospective study recruiting 2000 healthy volunteers by advertisement between February 2011 and July 2016 at the MRC London Institute of Medical Sciences. All subjects provided written informed consent for participation in the study, which was also approved by the National Research Ethics Service (09/H0707/69).

#### Image Acquisition

For both populations an equivalent CMR protocol was followed to assess LV structure and function using conventional two-dimensional retrospectively-gated cine imaging on a 1.5T magnet.^17, 59^ A contiguous stack of images in the LV short-axis plane from base to apex was used for volumetric analysis and trabecular phenotyping. Images were curated on open-source databases.^60, 61^

#### Image analysis

Segmentation of the short-axis cine images was performed using a fully convolutional network, a type of deep learning neural network, which predicts a pixelwise image segmentation by applying a number of convolutional filters onto each input image. The accuracy of image annotation using this algorithm is equivalent to expert human readers.^20^ Label maps were derived for all images in the cardiac cycle and LV volumes and mass were calculated according to standard guidelines,^62^ and then indexed to body surface area (BSA) using the Mosteller formula: 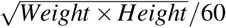, with weight in kg and height in cm.

Fractal dimension (FD) - a scale-invariant measure of trabecular complexity - was derived using a fully-automated algorithm executed from Matlab (Mathworks, Natick, MA) using custom-written code (autoFD, in GitHub repository at automated-fractal-analysis). Short-axis CMR images were pre-processed with bicubic interpolation to 0.25 mm x 0.25 mm pixels to enable consistent analysis between subjects acquired at different native resolutions. For each slice, a region of interest was defined within the midwall of the LV myocardium between the automated endocardial and epicardial segmentations. Subsequent image processing consisted of bias-field correction using histogram stretching, applying a region-based level-set algorithm and then binarization of the blood pool and myocardium.^63^ The trabecular borders were then detected using a Sobel filter and FD was calculated using a standard box-counting method in which the target image is overlain by a grid of known box size and the number of boxes containing non-zero image pixels is recorded. This process is repeated with box sizes between two pixels and 45% of the image size. Fractal dimension is defined as the negative gradient of an ordinary least-squares fit line to the logarithm of box size and box count. The FD values from all slices were interpolated using a Gaussian kernel local fit to a nine-slice template to allow comparison across subjects (see Figure 6 in the Supplement).

**Figure 6.**
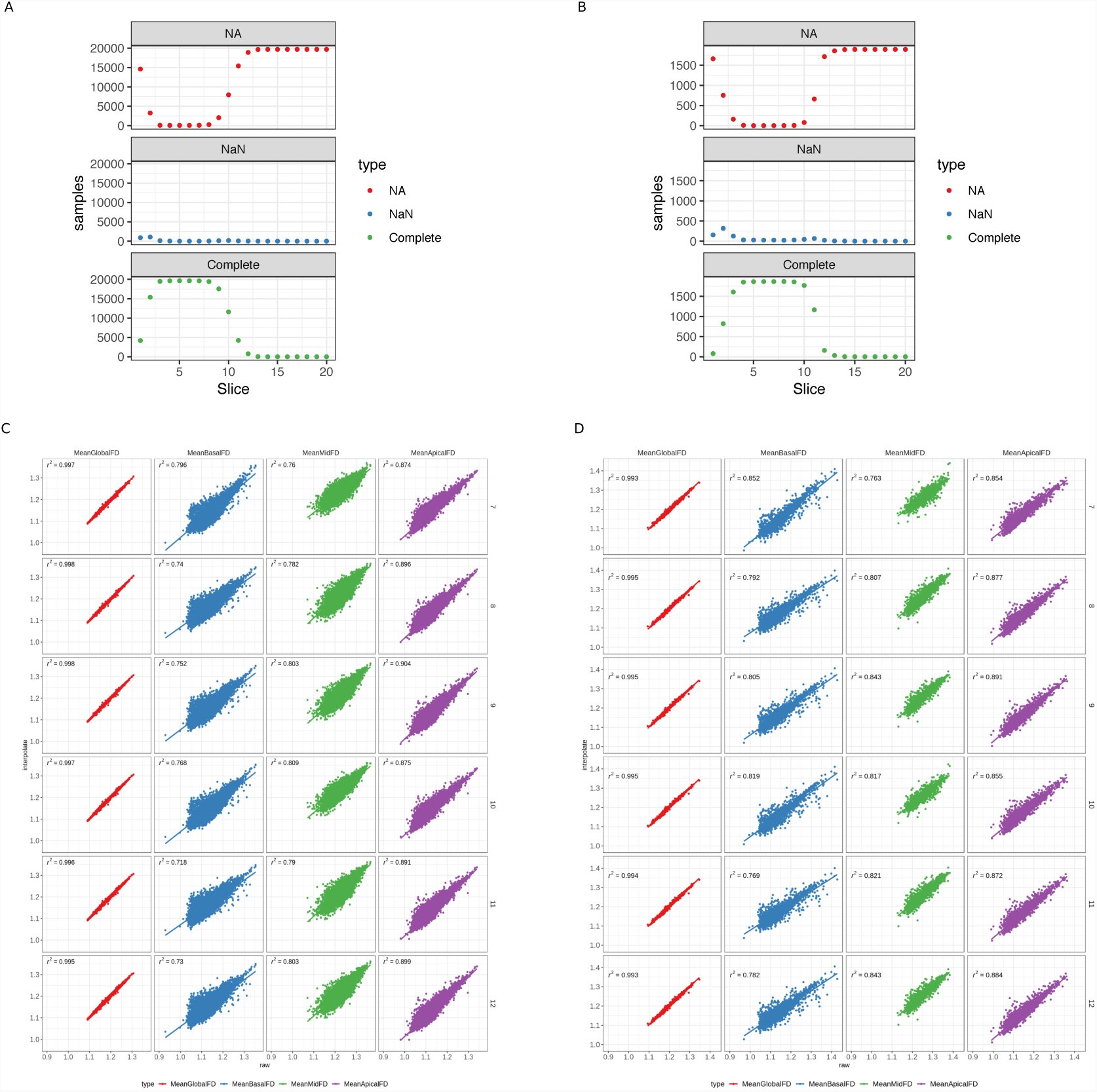
FD phenotypes. The upper panels show the distribution of CMR image slices where FD was successfully measured. Missing FD measurements per slice can either arise because a slice was not measured (NA) or the FD estimation failed due to poor image quality or estimated FD failing quality control (NaN). A. depicts the distribution in the UK Biobank samples, B. the distribution in the UK Digital Heart samples. The lower panels show the correlation between FD summary measures derived from the observed FD slice measurements and interpolated FD measurements per sample. Interpolated FD measurements per sample were derived by using a Gaussian kernel local fit to a different numbers of slice templates, allowing for direct slice comparisons across individuals. Different numbers of slices for interpolation were tested (rows). Columns show different summary measures, either mean FD across all measured slices or mean FD per slice region. C. depicts the correlation of the summary measures between measured and interpolated FD in the UK Biobank samples, D. the correlation in the UK Digital Heart samples.

For visual comparison of trabecular borders across individuals and genotypes, images were aligned by co-registering each ventricular segmentation to a common coordinate space and applying the same transformation to the corresponding greyscale image. For each slice, the trabecular and outer myocardial borders were extracted and their common center computed. The pixel positions of the edges were converted to radial coordinates (*θ, r*), with pre-defined step size of 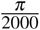. The outer myocardial border was then registered to a circle with radius one. The radial translocation at each angle required for the outer border registration was then applied to trabecular outline, making the outlines comparable across individuals (in GitHub repository at fractal-analysis-processing).

Pressure-volume loop analysis enables a detailed interpretation of cardiac physiology and ventricular work, but conventionally requires invasive catheterisation to obtain absolute pressure measurements which is not possible for population studies. Recently, a model-based framework combining non-invasive pressure measurements with CMR volumetry has been validated in a porcine model.^48^ Here we take advantage of consecutive CMR imaging and peripheral pulse-wave analysis (Vicorder, Wuerzburg, Germany) for dynamic volumetric analysis and central pressure estimation respectively, to non-invasively model left ventricular pressure-volume relationships throughout the cardiac cycle. Peak systolic pressure and maximal aortic distension on axial cine imaging were assumed to be synchronous allowing LV volume at peak-systolic pressure to be assessed. The indexed volume difference between end-diastole and end-systole over a single cycle was defined as stroke volume index, and over a minute as cardiac index. Indexed systemic vascular resistance was defined as the difference between mean arterial pressure and central venous pressure divided by cardiac index. In the absence of invasive catheter data, a value of 5mmHg for central venous pressure was assumed. LV diastolic pressures were assumed to be normal and rise during diastole from 4mmHg to 8mmHg at end-diastole.^64^ To understand the association of FD with pressure-volume dynamics, the mean ventricular FD value was associated with pressure and volume measurements during diastole and systole using linear regression, while controlling for contractility, systemic vascular resistance, trabecular mass and heart rate. Using this model, pressure-volume loops at comparable FD to the finite element model were plotted.

### Genotyping

#### Discovery cohort

UK Biobank genotypes release version 3 were used in the genetic association studies, for computing genetic principal components and for the Mendelian randomisation analyses. First, only unrelated or distantly related individuals were selected for further analysis based on the estimated identity of descent (IBD) provided by UK Biobank (via ‘ukbgene rel key.enc’). Selection of individuals from families is optimised to retain as many unrelated individuals as possible in the study i.e. in family trios only parents would be considered for analysis. For computation of genetic principal components, genotypes were formated into plink binary format,^65^ LD pruned (flag ‘–indep-pairwise 50kb 1 0.8’) and a minor allele frequency (MAF) threshold of 1% applied. The filtered dataset was used with flashpca version 2,^66^ to compute the first 50 principal components of the UK Biobank genotypes. Population substructures arising due to different ethnic origins of samples were examined by comparing the UK Biobank genotypes to genotypes from the HapMap Phase III study,^67^ for four ethnic populations (with subpopulations, Figure 7 in the Supplement). Download of reference data sets, fusion of UK Biobank and Hapmap data sets and PCA selection was done as described in plinkQC.^68^. Individuals that clustered with the main cluster of the PC1/PC2 plot, which was also the position of the main cluster of HapMap III individuals of European ancestry were kept for further analyses, as is standard in GWAS analysis to model a well mixed population. The genotypes of the remaining unrelated individuals were filtered for genetic variants that passed a MAF threshold of 0.1%. For association analysis this was achieved by providing a variant ID list with variant MAF > 0.001 to BGENIE.

**Figure 7.**
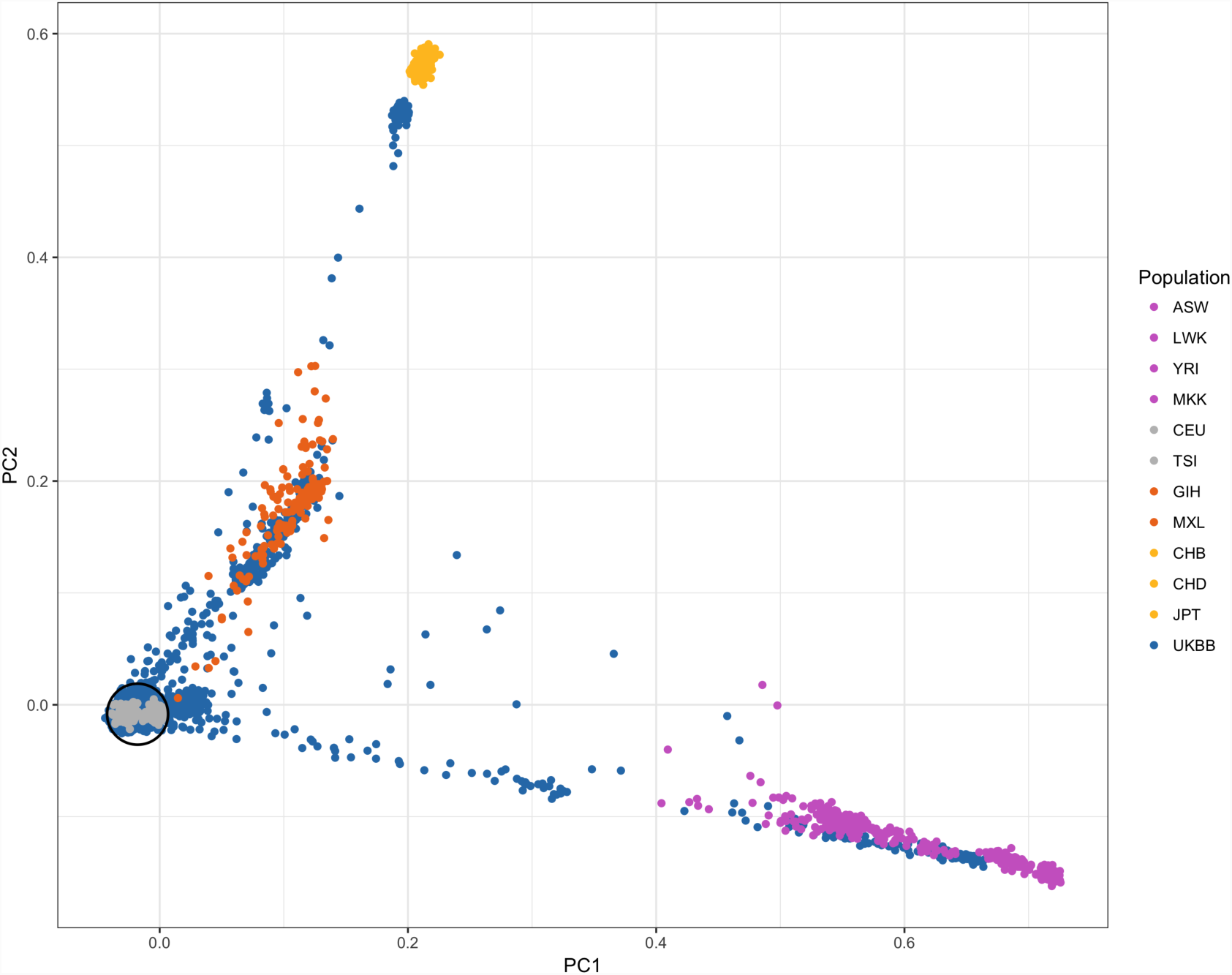
UK Biobank ethnicities. Principal components 1 and 2 of the principal component analysis on the combined genotypes of the UK Biobank and HapMap III datasets. UK Biobank individuals are depicted in blue, HapMap individuals colored by their ethnicity. UK Biobank individuals within 1.5 standard deviations distance from the center of the European HapMap individuals (grey) are selected for further analyses.

#### Validation cohort

UK Digital Heart project genotyping and genotype calling were carried out at the Genotyping and Microarray facility at the Wellcome Sanger Institute, UK and Duke-NUS Medical School, Singapore. Genotypes were assessed in five batches using Illumina HumanOmniExpress-12v1-1 (Sanger, two batches), Illumina HumanOmniExpress-24v1-0 (Duke-NUS, two batches) and Illumina HumanOmniExpress-24v1-1 chips (Duke-NUS). Genotypes were called via the GenCall software.^69^ For batches run on the same platform, genotype signals were combined and called in a single analysis, leading to three independent genotype batches. In order to avoid batch effects in genotype calling based on the probe sequences, probes targeting the same genotypes were checked for the concordance of the capture sequence. Genotyping probes common to all three platforms were selected and a common genotype dataset generated using PLINK v1.9,^65^ with plinkQC^68^ applied to assess the quality of the genotyping on a per-individual and per-marker level. In summary, the per-individual quality control included the identification of individuals with discordant sex information, missing genotype rates (more than 3% of genotypes not called) and heterozygosity rate outliers (three standard deviations outside of the mean heterozygosity rate). Population substructures arising due to different ethnic origins of samples were examined by comparing the sample genotypes to genotypes from the HapMap Phase III study,^67^ for four ethnic populations (with subpopulations, Figure 8 in the Supplement). Individuals that clustered with the main cluster of the PC1/PC2 plot, which was also the position of the main cluster of HapMap III individuals of European ancestry were kept for further analyses, as is standard in GWAS analysis to model a well mixed population. The per-marker quality control included filtering of genotypes with missing call rate in more than 1% of the samples and genotypes which significantly deviate from Hardy-Weinberg equilibrium (HWE, *p* < 0.001). After removing samples and genotypes that failed quality control, we confirmed that any pattern of missing genotype information was not batch-specific. To analyse these patterns, each pair-wise combination of batches was treated as a case-control set-up and the differential missingness of genotypes computed. Any genotype with significant differential missingness (*p* < 10^*-*5^) was removed from the dataset. After quality control, genotypes were phased and imputed to the combined 1000 Genomes^70^ and UK10K^71^ reference panel using SHAPEIT (version 2.r727)^72^ and IMPUTE2 (version 2.3.0).^73^ The window size for phasing was set to 2Mb, and the number of conditioning states per genotype to 200. The imputation interval was set to 3Mb, with a buffer region of 250kb on either side of the analysis interval. The effective population size was set to 20,000 and set the number of reference haplotypes to 1,000. For other non-specified parameters the default values were used.

**Figure 8.**
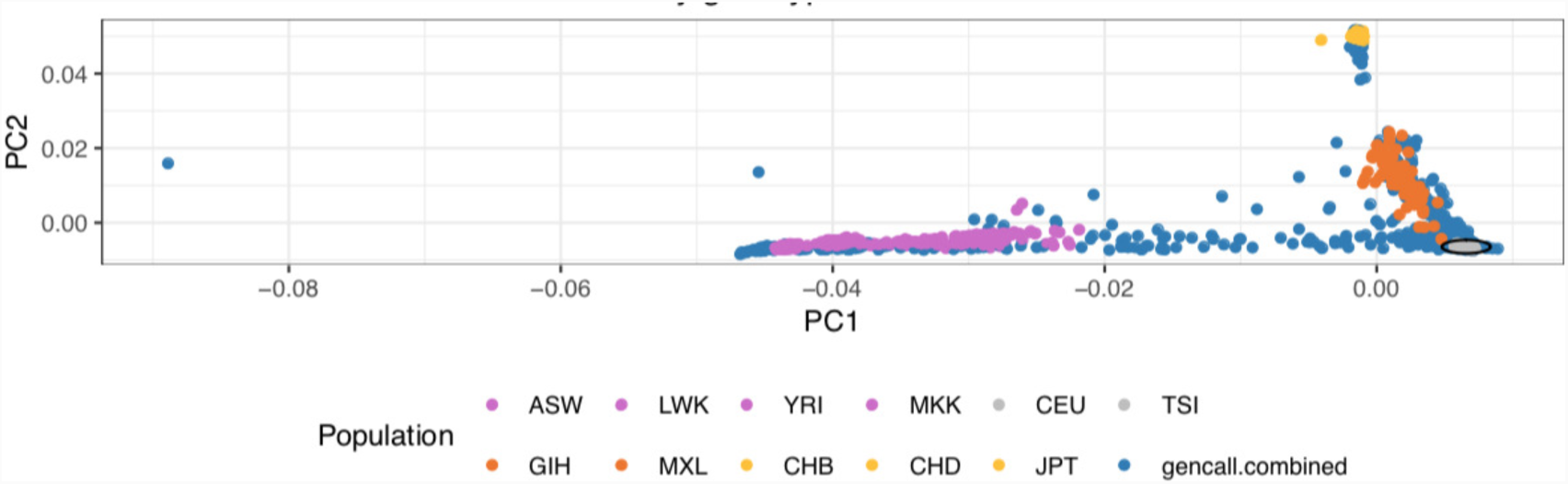
Digital heart study ethnicities. Principal components 1 and 2 of the principal component analysis on the combined genotypes of the UK Digital Heart study and HapMap III datasets. Digital Heart study individuals are depicted in blue, HapMap individuals colored by their ethnicity. Digital Heart study individuals within 1.5 standard deviations distance from the center of the European HapMap individuals (grey) are selected for further analyses.

### Association analysis

#### Univariate association

Genome-wide assocation studies (GWAS) for samples passing genotype and phenotype quality control (UK Biobank: 18,097, UK Digital Heart project: 1,129) were conducted using BGENIE v1.3 (https://jmarchini.org/bgenie).^16^ In the UK Biobank discovery cohort, we conducted univariate GWAS on the interpolated FD measurements per slice (9 independent GWAS) and the mean FD measurements per ventricular region (3 independent GWAS of basal, mid-ventricular and apical FD, Figure 1C), by fitting an additive model of association at each variant based on the genotype dosage of the imputed genotypes. In addition, we included sex, age, height, weight, BMI and principal components of the genotypes (PC1, 2, 6, 7, 14, 18, 21, 30, 47) as co-variates in the model (co-variates were determined by prior linear model fit of co-variates to phenotypes and selecting co-variates with p-value of effect size estimate < 0.05). The GWAS in the UK Digital Heart replication study used the same analysis setting and parameters as in the discovery cohort. To test for the effect of ventricular size, we conducted analogous GWAS in the discovery cohort, where we additionally included left ventricular end-diastolic volume as a co-variate. The univariate GWAS were adjusted for multiple-hypothesis testing by estimating the effective number of phenotype-association tests conducted^74^. The effective number of tests is estimated based on the eigenvalues **u** of the empirical trait-by-trait correlation matrix **C**. The effective number of tests is 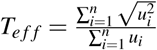, where *n* is the number of GWAS conducted i.e. *n* = 9 and *n* = 3 for per-slice and per-region GWAS, respectively.

#### Multi-trait meta-analysis

We used an approximate, multi-trait meta-analysis^75^ based on the univariate signed *t*-statistics of the 14,180,593 genetic variants for the nine per-slice FD GWAS. The multivariate test statistic is computed as 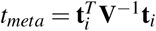, where **t**_*i*_ is the vector of the signed t-values of variant_*i*_ for the nine FD measurements, 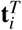 is a transpose of vector **t**_*i*_, **V**^*-*1^ is the inverse of the trait-by-trait correlation matrix **V**. **V**_*i, j*_ for each trait-trait pair is the correlation over the 14,180,593 estimated signed t-values of the two traits. *t*_*meta*_ is approximately chi-square distributed with 9 degrees of freedom and tests the null hypothesis that the genetic variant tested does not affect any of the nine traits.

#### Validation analysis

The effect size estimates of the FD GWAS of the discovery cohort (UK Biobank) and the FD GWAS of the validation cohort (UK Digital Heart project) were tested for concordance. For each locus in the discovery cohort that showed association with 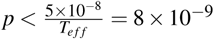, the genetic variant with the lowest p-value was selected. This selection was done for each univariate per-slice GWAS. The same loci-slice associations were selected in the validation cohort. Variants rs71394376 and rs71105784 are not present in UK Digital Heart genotypes and concordance was only tested at 14 out of 16 associated loci. The effect size estimated for the selected genetic variants in the discovery and validation data set were then compared for concordance, i.e. same effect size direction. To evaluate if the observed concordance was likely to arise by chance alone, an empirical p-value for concordance was estimated by randomly selecting slice-variant associations with a slice distribution as observed in the discovery associations. In each random selection step, the concordance of the observed and randomly selected slice effect size estimates was computed. The number of times the random concordance was greater or equal than the observed concordance was divided by the number of random selections (10,000) to yield the empirical p-value.

### Variant annotations

Variant annotations from previous studies were retrieved from Open targets genetics (https://genetics.opentargets.org/)^76^ for the PheWAS annotations, the GWAS catalogue^22^ for the GWAS annotations, the ENSEMBL regulatory build^45^ and GTEx v7 (https://gtexportal.org/home/) for the expression quantitative trait loci annotations. Annotations were reported when they passed the platform-specific significant thresholds (0.05 FDR on GTEx) or the commonly used GWAS threshold of 5 × 10^*-*8^.

### Functional enrichment analysis

We used GARFIELD version 2^46^ for functional enrichment analyses of genetic variants with multi-trait GWAS p-value < 10^*-*6^. The GARFIELD software package and pre-computed data for samples of Caucasian ancestry (LD and annotation data, minor allele frequencies of genetic variants and their distances to nearest transcription start site) were downloaded from https://www.ebi.ac.uk/birney-srv/GARFIELD.GARFIELD was run as described in the user manual at https://www.ebi.ac.uk/birney-srv/GARFIELD/documentation-v2/GARFIELD-v2.pdf. Annotation results (.perm GARFIELD output file) were filtered for input GWAS threshold (PThres < 10^*-*6^) and significance of enrichment (EmpPval < 10^*-*3^). Tissues of interest were selected (fetal heart, heart, fetal muscle, muscle, blood, blood vessel, epithelium) and all results of remaining tissues summarised in ‘Other tissues’. The full annotation results across all tissues can be found in Figure 14 in the Supplement).

#### Mendelian randomisation

Mendelian randomisation (MR) analysis was performed using all genetic variants with per-slice, univariate GWAS p-value 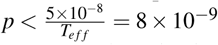. For variants associated with FD in multiple slices, only the slice association with the lowest p-value was used. All MR analyses were conducted using the R-package *TwoSampleMR* (https://github.com/MRCIEU/TwoSampleMR, version 0.4.18) which allows programmatic access to MRbase.^49^ MR analysis was performed using association results from independent studies (Two-sample MR). Traits of interest available on MRbase were stroke volume and QRS duration. The effect size of trabeculation on these traits was estimated with *TwoSampleMR* functions for weighted median, weighted mode, inverse variance weighting and MR-Egger. In addition, the Steiger test for directionality,^51^ leave-one-out sensitive analysis to determine the influence of single genetic variants on the overall effect, MR pleiotropy analyses and *I*^2^ analysis for assessing bias in MR-Egger analysis were conducted.^77^ For details refer to Mendelian randomisation in the Supplement.

#### Finite element modelling

We used a biomechanical model to assess the causative effect of varying trabecular morphology on cardiovascular physiology. To achieve this we compared ventricular behaviour with different degrees of trabecular complexity in the non-compact layer while keeping the total ventricular mass constant. A geometric model of the LV was represented by a symmetric ellipsoid truncated at two-thirds of the long axis to form a ventricular ‘base’. Ventricular dimensions were fitted to the median UK Biobank values for LV mass, end-diastolic volume and heart rate. Trabeculae were modelled as cylindrical strands orientated in the endocardial long-axis,^13, 78^ with total trabecular mass adjusted to the median value of UK Biobank subjects. Fractal dimension was calculated from cross-sections of the left ventricular model using the same methodology as for clinical imaging.

The ventricular model replicated *ex vivo* fibre orientations,^79^ and accounted for both active and passive material properties of the myocardium using a hyperelastic anisotropic constitutive framework. Specifically, the strain energy function *ψ* selected to model the cardiac tissue is 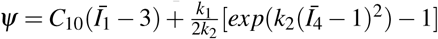, where *Ī*_1_ and *Ī*_4_ are the first and fourth invariant of the modified Cauchy-Green tensor 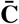 and *C*_10_, *k*_1_ and *k*_2_ are material parameters. Boundary conditions for the simulations included constraints on rigid motion and displacements of the ventricular base with implementation of a pre-load (defining left atrial pressure and inflow resistance) and after-load circuit (defining right atrial pressure, aortic/peripheral resistance, and capacitance). The models were discretized to allow finite element analyses with eight-node hexahedral elements into more than 10^4^ elements using Ansys Meshing (ANSYS Inc., Canonsburg, PA, USA). The property of differential fibre orientation within the ventricular wall, in which the sheets of muscle fibres describe a consistent helical pattern, was preserved by considering ventricular dynamics in nine separate myofibrillar sheets,^79, 80^ with linearly varying fibre orientation from +80 to −80 degrees with respect to the long-axis. Myocyte contraction in systole was simulated by changing the myocyte stiffness parameters to emulate the observed force/time data from cardiac fibres in response to intracellular calcium variation.^81^ Torsional motion was modelled by myocyte contraction patterns creating a realistic counter-clockwise apical rotation with respect to the base.^82–84^ The finite element problem was solved by means of the commercial code Abaqus (Abaqus 6.14, SIMULIA, Dessault Systemes). Simulations were performed under the assumption of quasi-static processes, so neglecting any inertia effects. Pressure-volume data are reported during steady state after a minimum of five simulated cardiac cycles. For details about the model parameters please refer to Table 14 and Table 15 in the Supplement.

**Table 14.** Material parameters in the computational model strain energy function.

| Material parameter | Diastolic Value | Maximum systolic value |
| --- | --- | --- |
| $C_{10}$ (kPa) | 0.2 | 7.2 |
| $k_1$ (kPa) | 1 | 180 |
| $k_2$ (-) | 2 | 2 |

**Table 15.** Resistances, compliance and atrial pressures in the *in silico* model pre-load and after-load circuits (Figure 4C).

| $P_{LA}$<br>(mmHg) | $R_1$<br>(mmHg*min/l) | $R_2$<br>(mmHg*min/l) | $R_3$<br>(mmHg*min/l) | $C$<br>(l/mmHg) | $P_{RA}$<br>(mmHg) |
| --- | --- | --- | --- | --- | --- |
| 5.25 | 0.2 | 0.375 | 18 | 0.0012 | 5 |

## Data availability

The genetic and phenotypic UK Biobank data are available upon application to the UK Biobank. The analysis code is freely available on GitHub (DOI:10.5281/zenodo.2571996).

## Supplementary Material

### Mendelian randomisation

Mendelian randomization (MR) studies can be thought of as randomised control trials where genetic variants are used as instrumental variables (IV) to infer the effect of an exposure variable on the outcome variable. The randomization is described as Mendelian based on Mendel’s laws of inheritance, i.e. the selection of alleles that an individual receives for a given genetic variant occurs at random during meiosis. This has two consequences, first the alleles are expected to be random with respect to confounders and second, they are causally upstream of the exposure traits. The effect of the exposure on outcome can then be inferred as the ratio of the genetic effect on the outcome over the genetic effect on the exposure.

In principle, there are two main types of MR analysis based on the study cohorts.^85^ In one-sample MR studies both the genetic variant-exposure and -outcome effect size estimates are obtained from the same cohort. In two-sample MR studies, these effect size estimates are obtained in independent cohorts. In times of large biobanks where thousands of individuals are measured for hundreds of traits, intermediate set-ups exist, where there is sample overlap between the cohort in which the exposure association was measured and the cohort of the outcome association.

#### MR assumptions and biases

The basis of MR studies is ‘vertical’ pleiotropy i.e. the genetic variants under investigation are associated with both traits considered because one trait is causal to the other trait. In addition, Mendelian randomisation studies make the following main assumptions:

- the instrument is associated with the exposure (IV1 assumption),
- the instrument only influences the outcome through exposure and not through any other pathway (IV2 assumption),
- the instrument is not associated with confounders (IV3 assumption).

However, there is a second type of pleiotropy, ‘horizontal’ pleiotropy, where the genetic variant is associated with the outcome through confounders or a pathway other than the exposure. ‘Horizontal’ pleiotropy is a violation of the IV2 and IV3 assumption and has to be addressed when conducting MR analysis as it can lead to incorrect inference of the causal effects.^86, 87^ In addition, one has to further consider the nature of ‘horizontal’ pleiotropy: if the mean effect of the ‘horizontal’ pleiotropy is zero, it is considered balanced and will not effect the effect size estimate of the causal effect (but can effect its standard error). If the mean effect is unequal to zero, it is considered directional and the extend of pleiotropy can be estimated (see MR Egger below).

If there is sample overlap between the exposure and outcome cohort, the overlap can lead to the correlation of the uncertainty in the genetic variant-exposure association and the genetic variant-outcome association. This can cause a bias of the causal effect estimates towards the confounded observational association (especially for weak instruments due to Winner’s curse^88^). IVs strongly correlated with the exposure (in practice defined as F statistic *>* 10) are less prone to this bias even in cohorts with overlapping samples.^89^

#### MR methods

A number of methods exist that can address violations to the IV assumptions outlined above. In the following, a selection of four methods used in this study are described in brief (for a comprehensive overview see^86–88^):

- **Inverse-variance weighted (IVW) linear regression.** If all IV are valid, IVW can be used to obtain an unbiased causal estimate of the exposure on the outcome. In IVW, the effect of the exposure on the outcome is estimated by linear regression of the effect size estimates of the genetic variant-exposure association on effect size estimates of the genetic variant-outcome association. The contribution of each IV to the overall effect is weighted by the inverse of the variance of the genetic variant-outcome effect and the intercept of the linear regression is constrained to pass through zero (no horizontal pleiotropy/balanced horizontal pleiotropy).^90^
- **MR Egger.** MR Egger works similar to IVW with the exception that the intercept is not restrained to pass through zero i.e. it allows to adjust for (unbalanced) pleiotropic effects. However, the effect estimates are only unbiased if the genetic variant-exposure associations and the pleiotropic effects are not correlated, i.e. the instrument strength is independent of direct effect (InSIDE assumption).^91^
- **Weighted median-based estimator.** The median-based estimator makes the assumption that the majority of IV are valid instruments. It is based on the ordered effect size estimate ratios for all IV-exposure to IV-outcome associations, weighted by standardised, inverse of the standard error of the Wald ratio estimated by the delta method (analogues to IVW). The weighted median-based estimator allows for unbalanced pleiotropy of the IVs and unlike MR Egger does not rely on the InSIDE assumption.^92^
- **Weighted mode-based estimator.** The mode-based estimator works based on cluster selection. It first clusters the IV into groups based on the similarity of the effect size estimates, and then selects the effect size estimate from the cluster with the largest number of IVs. The mode-based causal effect estimate is valid if the IVs in the largest cluster are valid.^49, 93^

In addition to these methods, there are a number of statistics that can be used to evaluate the validity of a given method. Both the IVW and MR Egger make the no measurement error (NOME) assumption, i.e. they consider the variance of the genetic variant-exposure association as negligible.^90^ In IVW, the presence of large measurement error (violation of NOME) can lead to weak instrument bias. The strength of the instruments can be assessed with the F-statistic and as above for overlapping sample sizes, IVs which strongly correlated with the exposure (in practice defined as F statistic *>* 10) are less prone to this bias.^94^ However, the F statistic is only a proxy for the true, but unknown parameter of interest, the F parameter. In addition to computing the cohort F statistic, one can also estimate a lower bound on the true F parameter as described in [95, Appendix A3]. For MR Egger and NOME violation, the causal effect size estimates can suffer from regression dilution bias which attenuates the causal effect estimate towards the null,^77^ The MR Egger *I*^2^ statistic can be calculated to evaluate the magnitude of dilution bias, e.g. an *I*^2^ of 90% will lead to a 10% underestimation of the effect size.^77^ The mode-based estimator can be implemented with or without the assumption of NOME.^49, 93^

MR analysis can be used to infer whether there is a causality between exposure and outcome and which direction the effect takes i.e. exposure-outcome or outcome-exposure. With the MR Steiger test, the directionality can be assessed based in the absolute correlations of the genetic variants with the exposure and outcome (with Steiger’s Z-test for correlated correlations within a population). Based on the Z statistic and the associated Steiger p-value one can then test at a predefined level *α* if one accepts the causal association for the model.^51^

**Figure 9.**
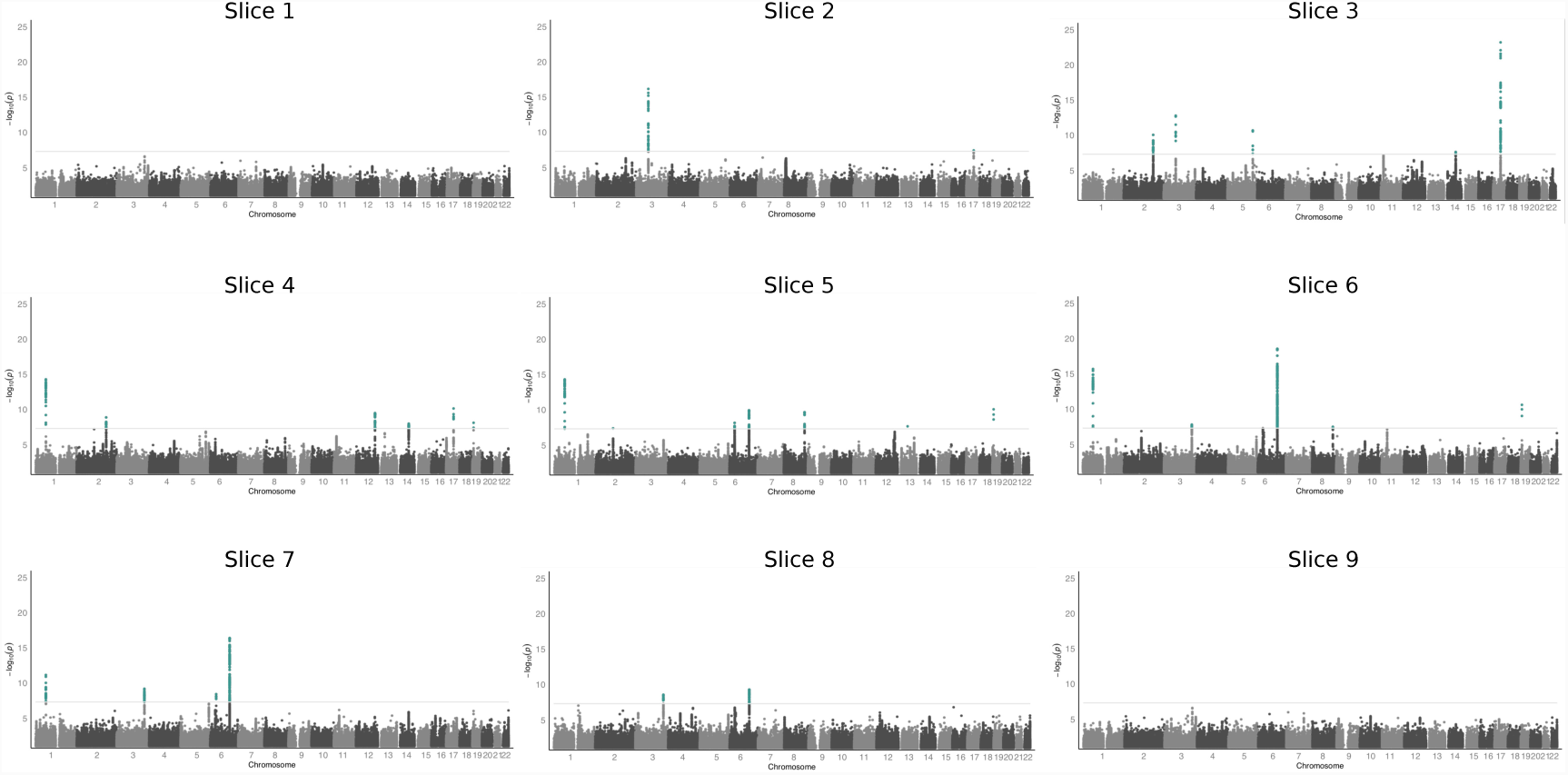
Per-slice FD GWAS results. Manhattan plots of the independently conducted, nine univariate GWAS on the FD measurements per slice. The p-values were multiplied by the effect number of independent phenotypic tests *T*_*e f f*_ = 6.6. The horizontal grey line is drawn at the level of genome-wide significance: *p* = 5 × 10^*-*8^.

**Figure 10.**
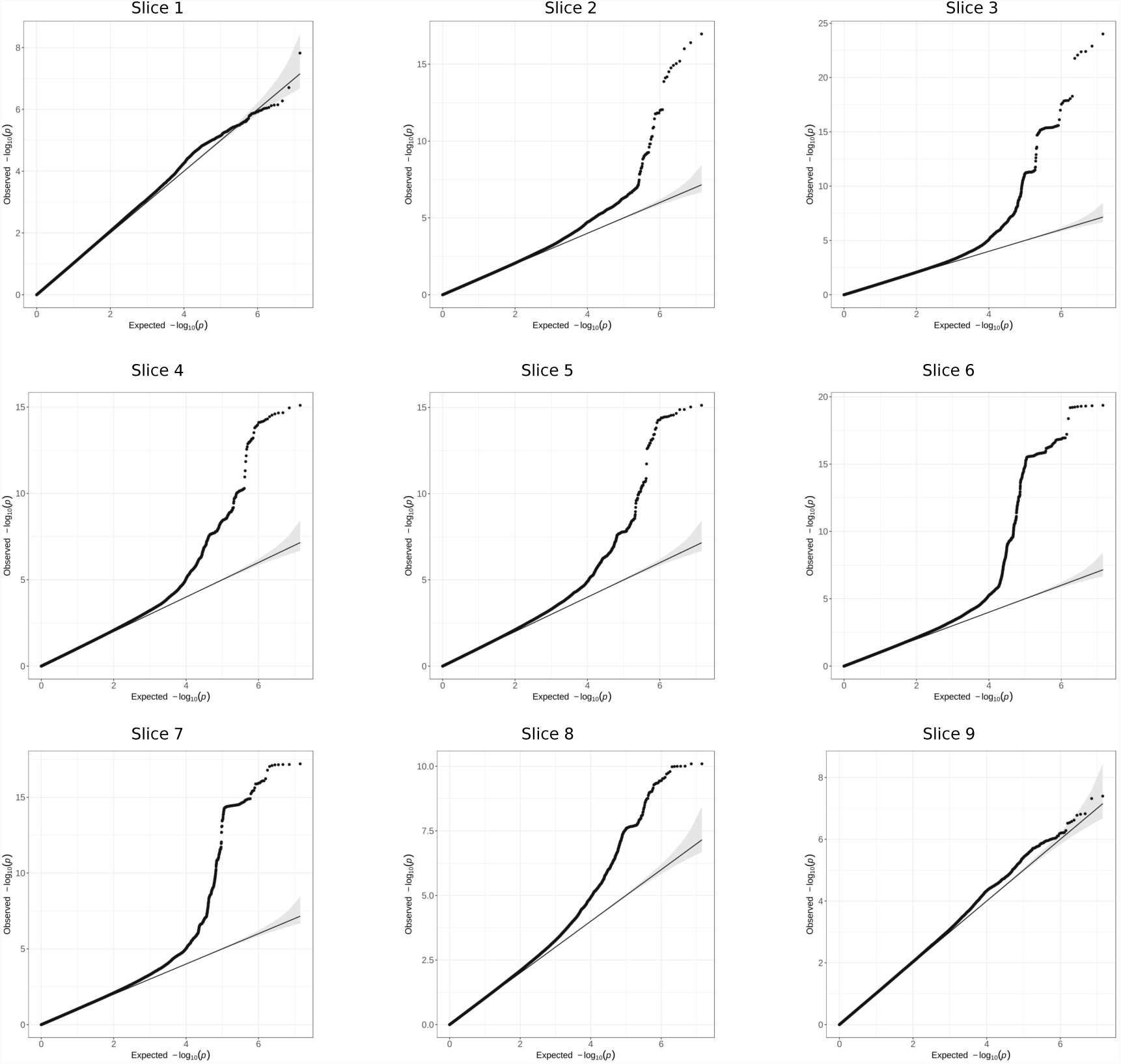
Per-slice FD GWAS calibration. Quantile-quantile plots of the independently conducted, nine uni-variate GWAS on the FD measurements per slice. The observed genome-wide p-values are plotted against equally spaced values in [0, 1] of the same sample size (expected p-values). The diagonal line starts at the origin and has slope one.

**Figure 11.**
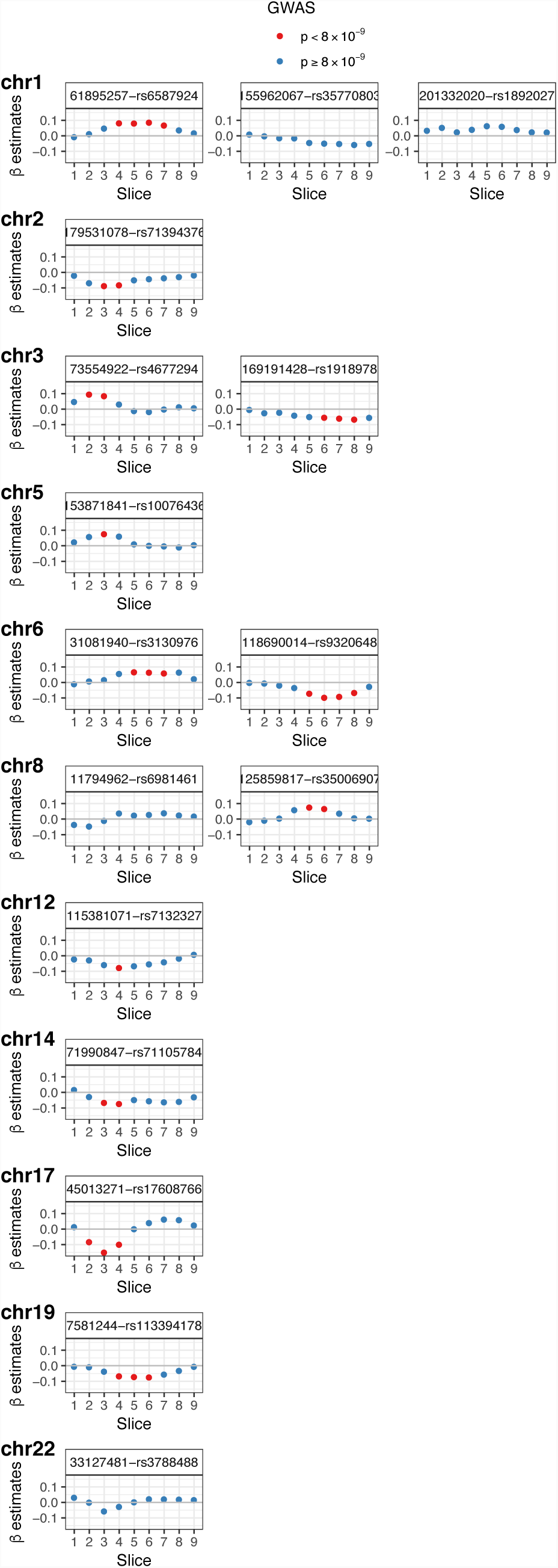
Distribution of effect size estimates. Effect size distribution of loci with genetic variant associations of 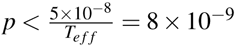 in any uni-variate per-slice FD GWAS. Distribution shown for each locus (indicated by chromosomal position and lead genetic variant in subplot title) across all slices and effect size colour-coded by p-value of the association. Effect sizes of association with *p* < 8 × 10^*-*9^ is similar in adjacent slices. Variants with no *p* < 8 × 10^*-*9^ in the univariate per-slice FD GWAS (all blue) were discovered in the multi-trait meta-analyses.

**Figure 12.**
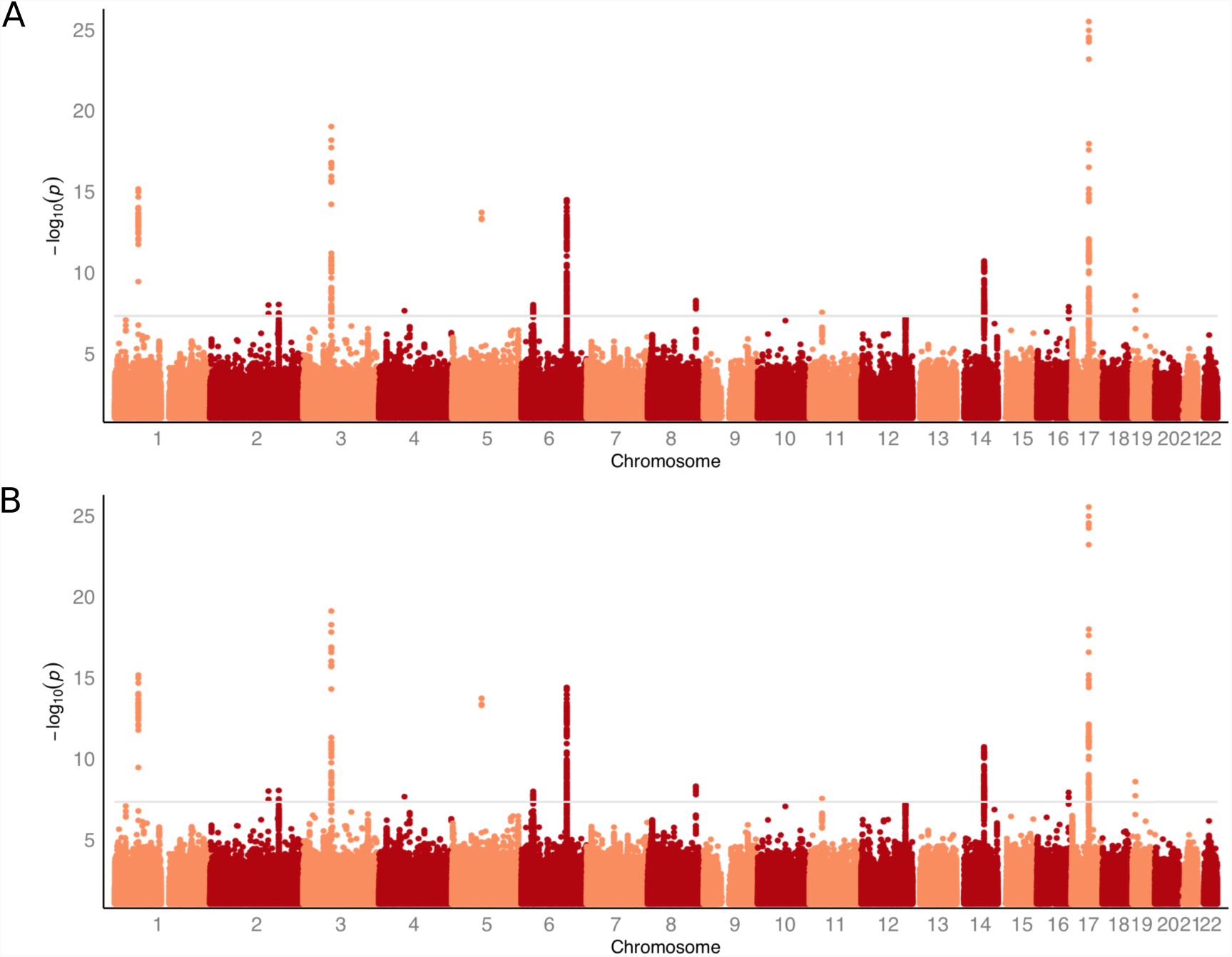
Comparison of FD GWAS with end-diastolic volume as co-variate. A. Manhattan plot based on meta-analysis GWAS with end-diastolic volume of the left ventricle as co-variate and B. Manhattan plot based on meta-analysis GWAS without end-diastolic volume as co-variate (same as Figure 2B; shown for comparison). Other co-variates and analysis parameters (as described in methods) were kept the same in A and B. The horizontal grey lines are drawn at the level of genome-wide significance: *p* = 5 × 10^*-*8^.

**Figure 13.**
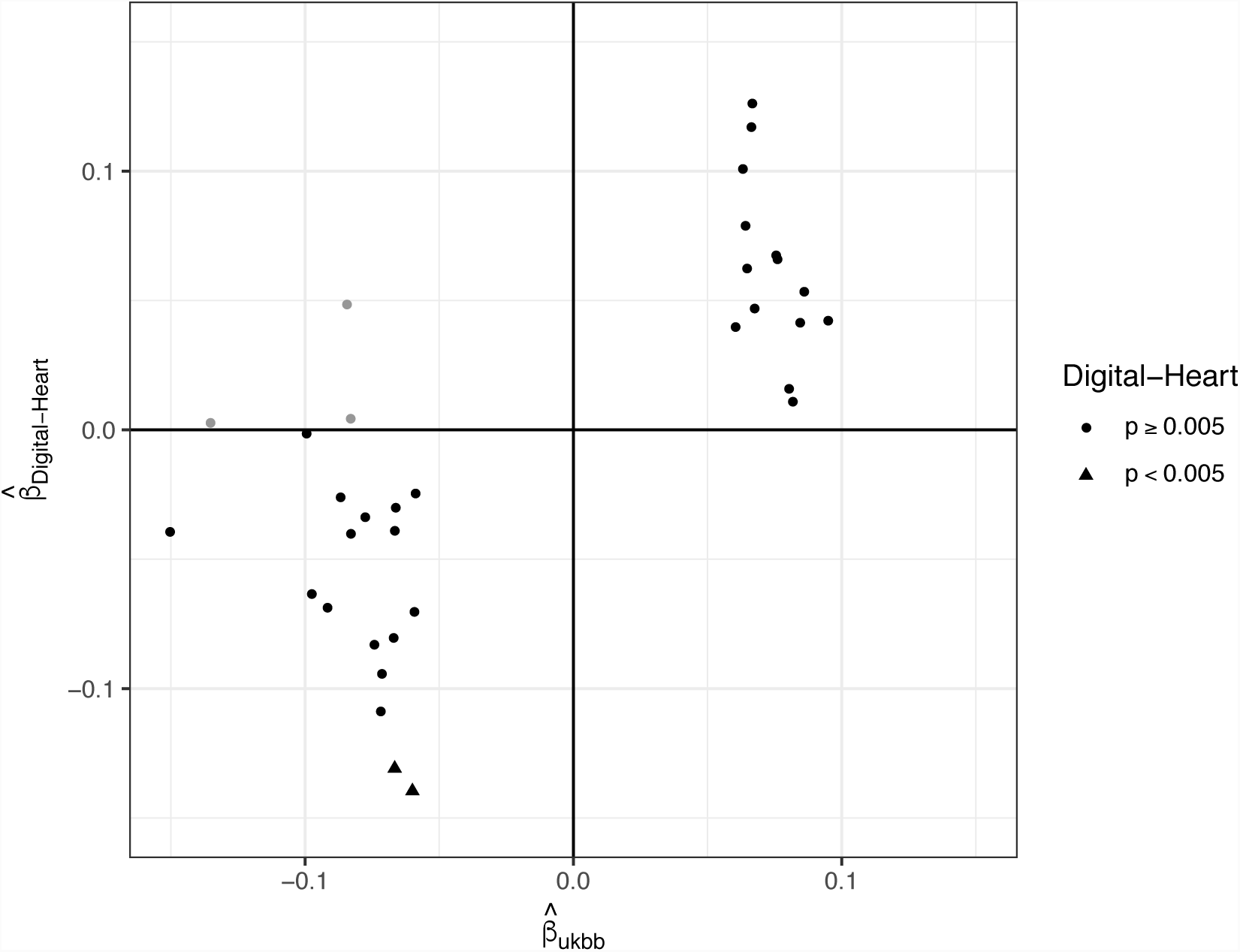
Effect size estimate concordance in discovery and validation cohort. For each of the nine uni-variate, per-slice FD GWAS, the effect size estimates of the genetic variants with the smallest p-value for each of the independent loci in the discovery cohort (ukbb) were selected. For some variants, associations passing the GWAS threshold of 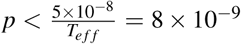 were discovered in more than one of the nine uni-variate GWAS FD slices; for these variants all effect size estimates were selected. These estimates were plotted against the corresponding slice-variant associations in the validation GWAS (Digital-heart). Non-concordant estimate pairs are depicted in light grey. Effect size estimates passing the validation p-value threshold of *p* < 0.005 are depicted as triangles.

**Figure 14.**
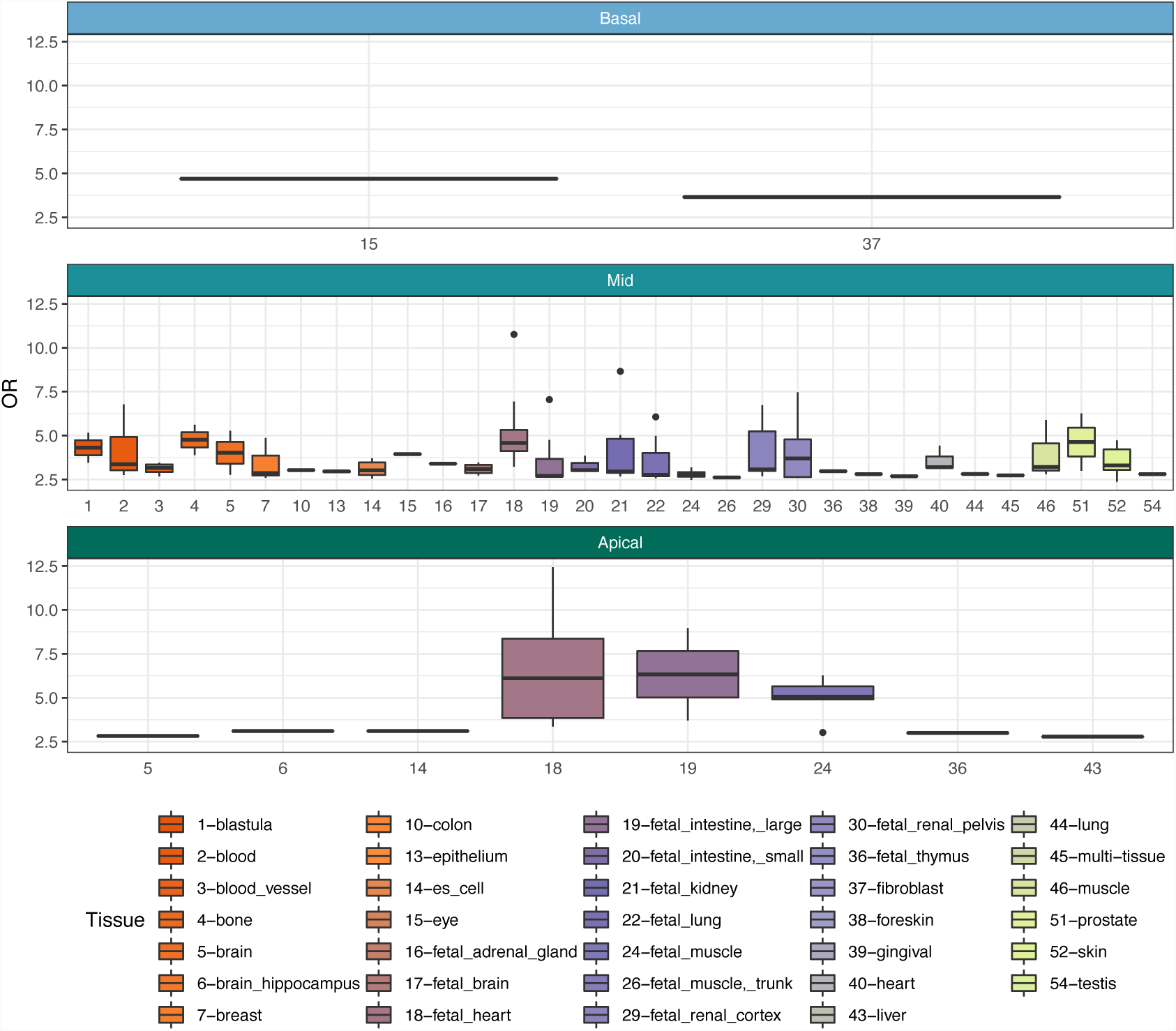
Enrichment of trabeculation associated variants in DNaseI Hypersensitive sites for all available tissues in GARFIELD. GARFIELD was used to compute the functional enrichment (odds ratio, OR) of genetic variants associated with the trabeculation phenotypes (*p* < 10^*-*6^) for open chromatin regions. Enrichment analyses were done for each cardiac region independently and results across all available studies per tissue are depicted in boxplots.

**Figure 15.**
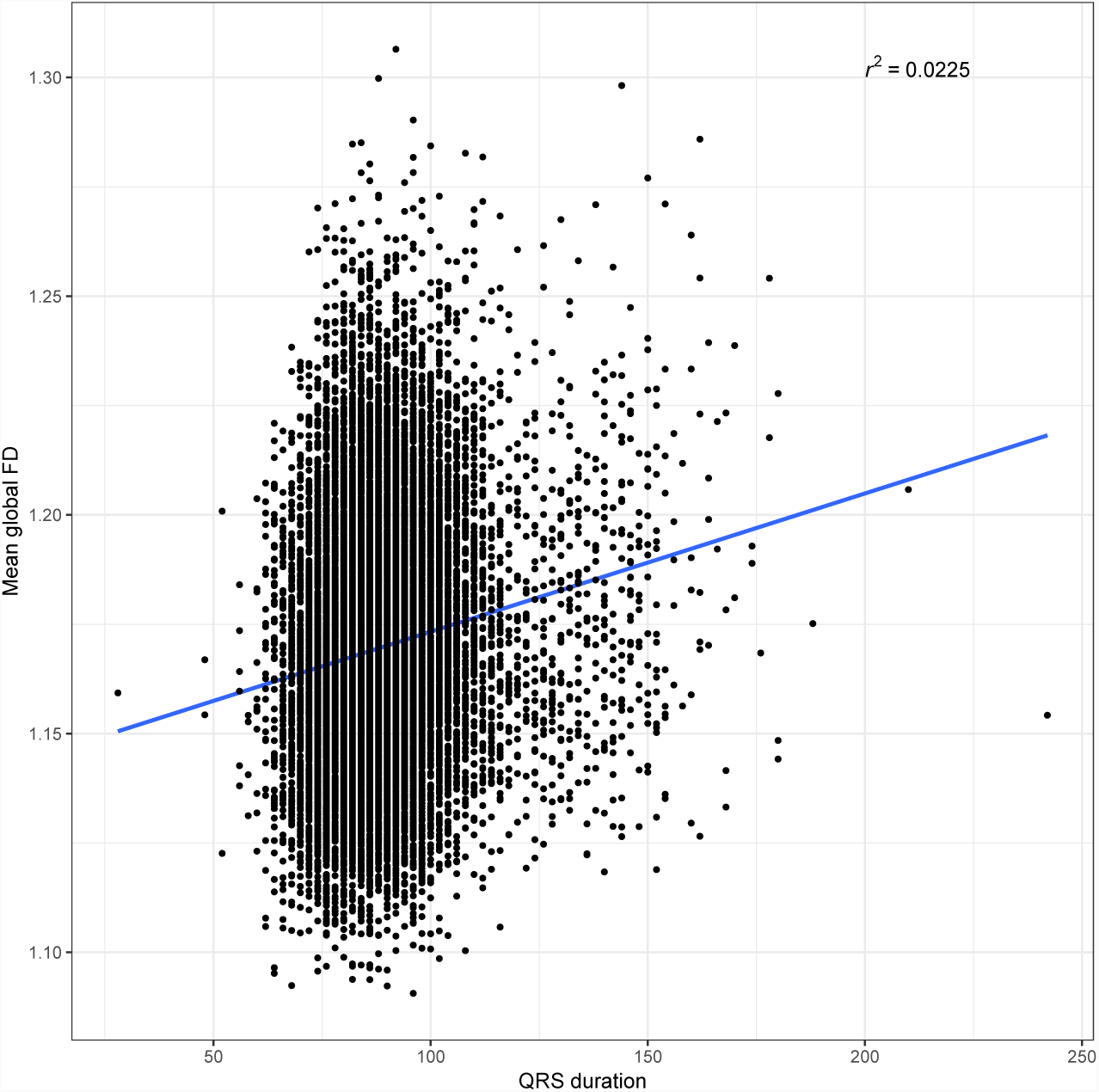
Pearson correlation of global FD and QRS duration. QRS duration phenotype from UKB ID: qrs duration f12340 2 0. The Pearson correlation coefficient is indicated in the upper right corner.

**Figure 16.**
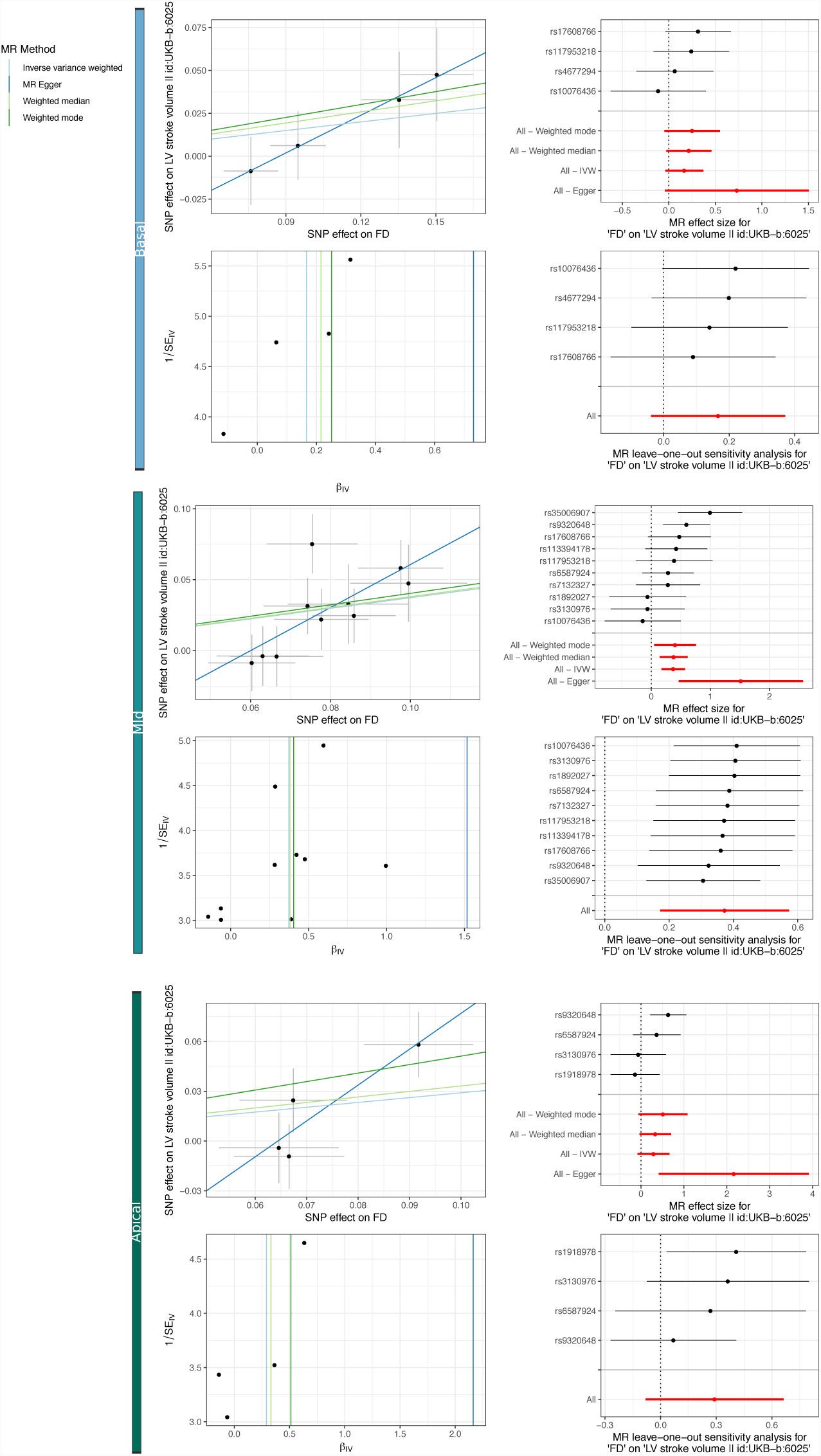
MR analysis of trabeculation on stroke volume. For each ventricular region (basal, mid-ventricular and apical; color-coded column strips) four panels showing a scatter plot (upper left), forest plot (upper right), funnel plot (lower left) and leave-one-out sensitivity plot (lower left). Scatter plots depict the genetic variant-exposure effect versus the genetic variant-outcome effect. Forest plots show the contribution of each genetic variant to the overall estimate (black) and combined as a single genetic instrument (red) for the four tested MR methods (see legend). Funnel plots depict the instrument strength against the causal effect of each instrument as a single IV. Vertical lines indicate the average estimated effect for the tested MR methods. Strong instruments are close to the estimated average effect, while weak instruments spread evenly on both sides. Leave-one-out plots show the results of MR analysis (IVW only) where each genetic variant is sequentially excluded and can indicate if there are any single variants that drive the MR results.

**Figure 17.**
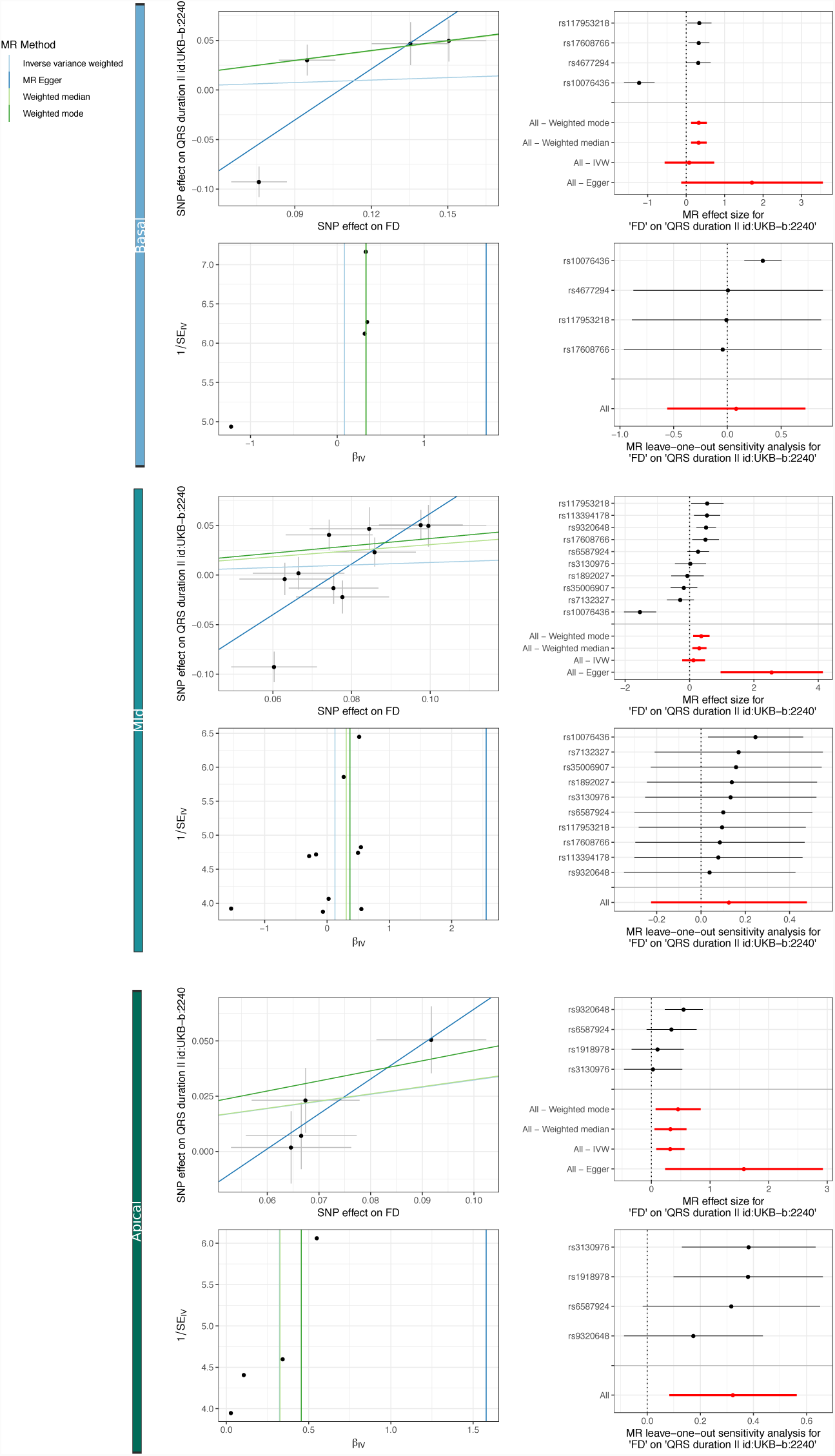
MR analysis of trabeculation on QRS duration. For each ventricular region (basal, mid-ventricular and apical; color-coded column strips) four panels showing a scatter plot (upper left), forest plot (upper right), funnel plot (lower left) and leave-one-out sensitivity plot (lower left). Scatter plots depict the genetic variant-exposure effect versus the genetic variant-outcome effect. Forest plots show the contribution of each genetic variant to the overall estimate (black) and combined as a single genetic instrument (red) for the four tested MR methods (see legend). Funnel plots depict the instrument strength against the causal effect of each instrument as a single IV. Vertical lines indicate the average estimated effect for the tested MR methods. Strong instruments are close to the estimated average effect, while weak instruments spread evenly on both sides. Leave-one-out plots show the results of MR analysis (IVW only) where each genetic variant is sequentially excluded and can indicate if there are any single variants that drive the MR results.

## Acknowledgments

The research was supported by the British Heart Foundation (NH/17/1/32725); the National Institute for Health Research Biomedical Research Centre based at Imperial College Healthcare NHS Trust and Imperial College London; and the Medical Research Council, UK. P.M.M. also has been in receipt of generous personal and research support from the Edmond J Safra Foundation and Lily Safra, an NIHR Senior Investigator’s Award, the Medical Research Council and the UK Dementia Research Institute. The authors would like to thank Dr Hideaki Suzuki, previously of Department of Medicine, Imperial College London, for his work on pre-processing the image data, and Prof Roberto Fumero, at the Politecnico di Milano, Italy, for advice on the finite element modelling. The authors would also like Virginie Uhlmann (EMBL-EBI) for advice on radial image registration and George Davey Smith (University of Bristol) for advice on Mendelian randomisation analysis

## Author contributions

H.V.M. and T.J.W.D. performed the formal analysis and co-wrote the manuscript; M.S. and M.L.C. performed the *in silico* modelling; T.J.W.D and A.M. collected and analysed image data; W.B., P.T., J.C. and D.R. developed the computational phenotyping; P.M.M., E.B., S.A.C. and D.P.O. provided critical interpretation of the results; E.B., S.A.C. and D.P.O. conceived the study, managed the project and revised the manuscript. All authors reviewed the final manuscript.

